# Linking species coexistence mechanisms to biodiversity effects on productivity in multispecies communities under environmental variation

**DOI:** 10.64898/2026.11.26.768821

**Authors:** Xiangyu Liu, Oscar Godoy, Caroline Daniel, Noémie A. Pichon, Seraina L. Cappelli, Eric Allan

**Affiliations:** Institute of Plant Sciences (IPS), Bern University, Bern, Switzerland; Estación Biológica de Doñana (EBD-CSIC), Sevilla, Spain; Institute of Ecology and Evolution, University of Bern, Bern, Switzerland; Swiss Federal Research Institute WSL, Birmensdorf, Switzerland; Swiss Federal Research Institute WSL, Davos Dorf, Switzerland; Oeschger Centre for Climate Change Research, University of Bern, Bern, Switzerland; Centre for Development and Environment, University of Bern, Bern, Switzerland

## Abstract

Linking coexistence mechanisms and biodiversity effects on ecosystem functioning could provide mechanistic understanding of biodiversity change. However, this framework remains poorly integrated with functional traits and rarely applied to multispecies communities under varying environmental conditions. Here, we coupled field-parameterized models of interactions between 18 perennial plant species, with data from a biodiversity-functioning experiment manipulating species richness, nitrogen, and functional composition based on fast-slow traits. We connected coexistence mechanisms (niche differences, competitive ability differences and the contribution of indirect interactions to coexistence) to biodiversity effects on productivity (complementarity, intra-and interspecific selection effects) within each community. When facilitation was excluded, complementarity was higher in slow-growing, niche differentiated communities and interspecific selection (dominance) effects were higher in communities with large competitive ability differences. Coexistence mechanisms explained part of the relationships between traits and biodiversity effects but direct trait effects were stronger and varied with nitrogen and season. Importantly, including facilitation disrupted these links by promoting the coexistence of low-biomass, fast-growing species that contributed least to complementarity. Our results show when independently estimated coexistence mechanisms explain biodiversity effects on productivity, and why greater opportunities for coexistence created by facilitation need not translate into higher complementarity.

## Main

Ecologists have long recognized that biodiversity enhances ecosystem functioning^1–3^. Diverse communities function better either because species are more niche differentiated, leading to greater complementarity^4,5^, or because high-performing species competitively dominated communities and contribute disproportionately to functioning through selection (dominance) effects^6,7^. These interpretations implicitly link biodiversity effects to coexistence mechanisms: complementarity is expected to arise from niche differences, whereas selection effects reflect competitive ability differences between species^8,9^. However, a clear link is not inevitable as theoretical models show complementarity effects can arise even in the absence of niche differences^10^ and complementarity effects are often maximized under even abundances, which are not guaranteed under stable coexistence^11,12^. In fact, such links have rarely been empirically demonstrated, particularly in multispecies communities using independent measurements of species interactions and biodiversity effects^10,13–15^. Further, to use coexistence theory to predict global change effects on diverse communities, we need to understand how links between coexistence and biodiversity effects depend on functional traits, species interaction types and environmental context (Fig. 1).

**Fig. 1.**
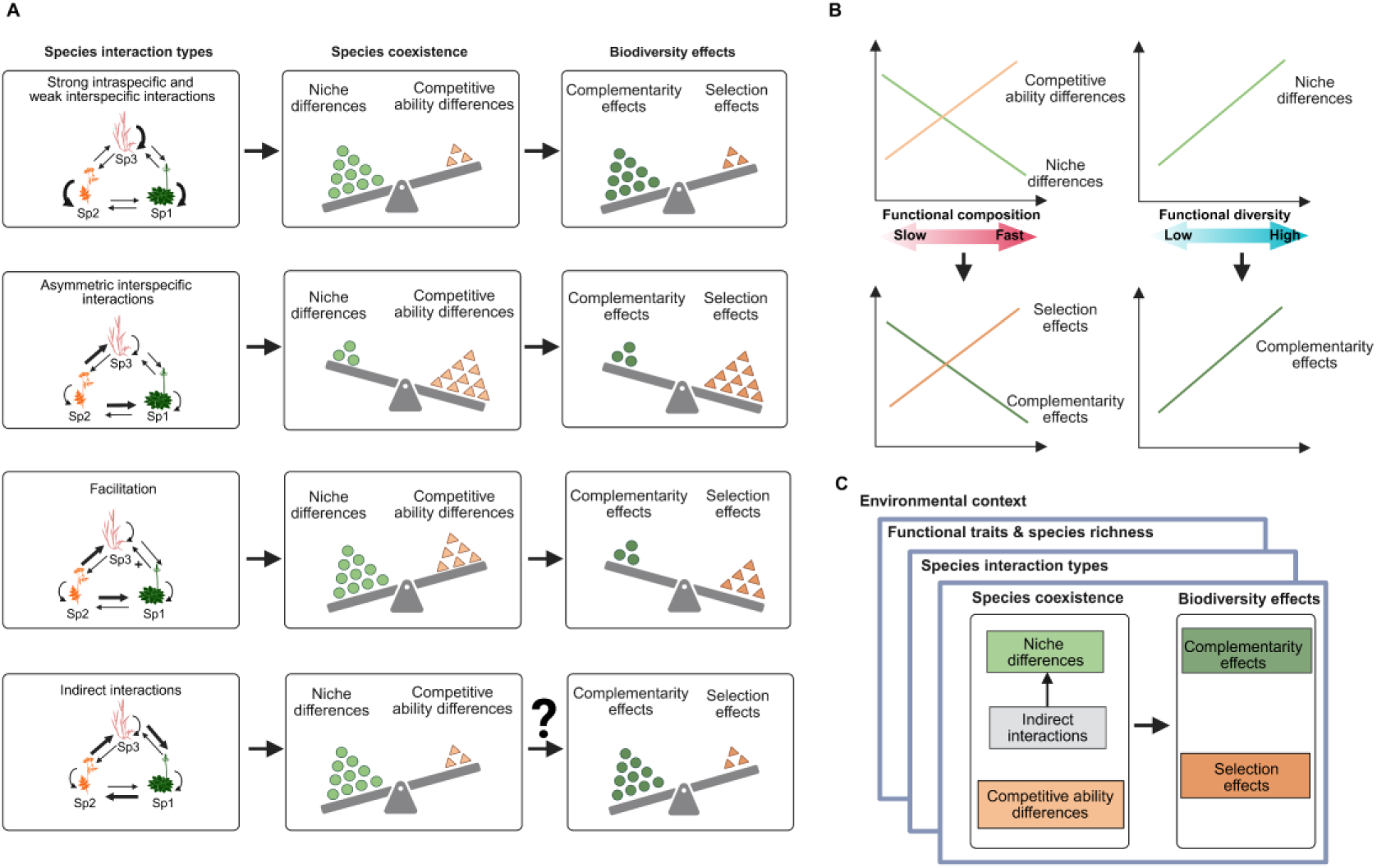
Coexistence mechanisms and biodiversity effects emerging from different types of species interactions (A). In the classical coexistence framework^21,22^, coexistence is promoted by stabilizing niche differences and by equalizing processes that decrease competitive abilities differences. Strong intraspecific and weak interspecific interactions (arrow width reflects interaction strength) illustrate stabilizing niche differences that can promote coexistence despite competitive ability differences. We hypothesize that this stabilization is associated with higher complementarity effects and lower interspecific selection (dominance) effects. In contrast, asymmetric interspecific interactions illustrate stronger competitive ability differences that can favor dominance and exclusion when stabilization is insufficient to offset competitive ability differences. We hypothesize that these conditions are associated with higher interspecific selection effects and lower complementarity effects. Facilitation (indicated by a black arrow denoted with +) can also promote coexistence by expanding niche differences but may allow facilitated species to persist only at low abundances, thus contributing little to complementarity. Indirect interactions (e.g., the rock-paper-scissors intransitive competition loop) can enlarge the feasibility domain (the structural analogue of niche differences) and create additional pathways for coexistence^25^ but their effects on biodiversity effects remain unclear. (B) Functional traits influence biodiversity effects by modifying coexistence mechanisms. Slow-and fast-growing species differ in resource economic strategies and in their adaptation to low-versus high-resource environments^18^. Slow-growing species may exhibit greater niche differences and complementarity, whereas fast-growing species often share similar light acquisition strategies, leading to stronger competitive ability differences and larger interspecific selection effects. Variance in functional traits (functional diversity) can promote niche differences and complementarity when species different in resource use strategies. However, whether trait effects on biodiversity effects can be explained by their impacts on coexistence mechanisms remains unknown. (C) Environmental context (e.g., nitrogen availability and foliar pathogens), species richness and functional traits (i.e., functional composition and functional diversity) and species interaction types may reshape the relationships between coexistence mechanisms and biodiversity effects. Intraspecific selection effects are excluded, and interspecific selection effects are labelled as selection effects for simplicity, but we would expect that high complementarity is associated with strongly negative intraspecific selection effects and therefore that niche differences should lead to negative intraspecific selection.

Global change may alter links between coexistence mechanisms and biodiversity effects by shifting functional trait composition. Land use intensification, for example, often shifts communities from slow-growing, resource-conservative species towards fast-growing, resource-acquisitive species^16,17^. Such changes could impact both coexistence and biodiversity-functioning mechanisms. Functional diversity is thought to promote niche differences and complementarity, whereas functional composition (e.g., whether communities are dominated by fast-or slow-growing species) should influence competitive ability differences and selection effects (Fig. 1B)^18–20^. Further, slow-growing species may exhibit greater differentiation in nutrient competition, potentially leading to larger niche differences. However, coexistence mechanisms depend on long-term population outcomes while biodiversity effects are estimated from short-term biomass production. This means factors that alter species relative abundances without affecting long-term persistence may alter biodiversity effects, but not the probability of coexistence. It therefore remains unclear whether trait-driven biodiversity effects can be explained by trait effects on coexistence mechanisms.

The relationship between coexistence and biodiversity effects depends on the strength and direction of species interactions, and whether interactions are direct or indirect. Stabilizing niche differences arise when species limit themselves more strongly than they limit one another (Fig. 1A, strong intraspecific and weak interspecific competition), whereas competitively ability differences favor some species over the others (Fig. 1A, strong asymmetric interspecific competition)^21,22^. Coexistence is promoted when niche differences are sufficient to offset competitive ability differences. However, facilitation and indirect interactions can provide additional opportunities for coexistence in multispecies communities^23–25^. Facilitation occurs when one species benefits another, for example by buffering temperature extremes and increasing soil moisture^26–28^. This positive interaction can promote coexistence and might also increase complementarity in multispecies communities^26^. However, when facilitation allows species to persist only at low abundances, it could promote coexistence without increasing complementarity^29^, thereby decoupling the links between coexistence and biodiversity effects (Fig. 1A). Indirect interactions occur when one species alters another by changing the abundance of a third species^30^, such interaction chains can create conditions for coexistence that do not occur with purely pairwise mechanisms (i.e., Fig. 1A, rock-paper-scissors intransitive competition loops). There is no theory or empirical data on whether biodiversity effects, like complementarity effects, are higher in communities where indirect interactions expand opportunities for species to coexist. Recent theoretical advances have made it possible to quantify coexistence mechanisms, including niche and competitive ability differences, while accounting for facilitation and indirect interactions^31,32^. This allows us to test whether the additional coexistence opportunities generated by facilitation and indirect interactions also result in higher functioning.

Global change drivers frequently alter resource availability, particularly nitrogen, and pressure from key plant enemies, like foliar pathogens^33–35^. Both factors are likely to change coexistence mechanisms and biodiversity effects on ecosystem functioning^8,36,37^. In grasslands, nitrogen enrichment typically reduces niche differences and increases competitive asymmetry, weakening coexistence and shifting diversity effects from complementarity towards selection^38–42^. Similarly, foliar pathogen removal can erode coexistence by weakening negative intraspecific density, in turn, potentially reducing complementarity effects^43^. Reduction in foliar pathogens may also alter competitive hierarchies, if some species benefit more from enemy release than others^44–47^, thereby changing selection effects. A predictive framework unifying coexistence mechanisms and diversity effects under global change will require a better understanding of how resources, natural enemies and traits shape their interrelationships (Fig. 1C).

In this study, we test how coexistence mechanisms in multispecies communities predict ecosystem functioning across environmental and functional trait contexts. We integrated two independent datasets from a long-term grassland biodiversity experiment, the PaNDiv experiment. First, we quantified biodiversity effects on biomass production for communities differing in plant species richness, functional diversity and composition, nitrogen addition and foliar pathogen abundance^48–50^. We used Fox’s tripartite partitioning to further decompose the selection effects of Loreau and Hector into two components^51,52^, which we refer to here as interspecific selection effects (dominance effects sensu Fox) and intraspecific selection effects (trait-dependent complementarity effects sensu Fox)^20,52^. We used these terms, instead of Fox’s terminology, to highlight the fact that the selection effects have been partitioned into two components, while the complementarity effect is the same in both partitions^51,52^. Intraspecific selection effects are typically negative and occur when low performing species increase in mixture without affecting other species^20,52^. This could indicate strong negative density dependence for low biomass species and might therefore link to niche differences^52^. In contrast, interspecific selection effects would be expected to be positive if high biomass species competitively dominate mixtures, so these would be expected to link to competitive ability differences. Second, we used an independent phytometer experiment, conducted in the field, to estimate species interactions and intrinsic growth rates for the species present in the PaNDiv experiment, under nitrogen and fungicide treatments^53^. We then used the structuralist approach to calculate coexistence mechanisms, including niche differences (derived from the size of the feasibility domain, a metric accounts for all sources of variation promoting niche differences), competitive ability differences and the contribution of indirect interactions for each multispecies plot (based on the sown species composition) with and without facilitation^31,32,54^. Finally, we used structural equation models to test links between coexistence mechanisms and biodiversity effects across environmental and functional trait contexts. The combination of these two independent datasets into a single analysis constitutes a hard test of the role of biotic interactions in driving ecosystem functioning.

We test four hypotheses: (1) Communities with greater niche differences will have higher complementarity effects and more negative intraspecific selection effects, while those with higher competitive ability differences will have larger interspecific (dominance) selection effects (Fig. 1A); (2) functionally diverse communities, and communities dominated by slow species, will have higher niche differences and complementarity effects, while fast dominated communities will show greater competitive ability differences and interspecific selection (dominance) effects (Fig. 1B); (3) Facilitation and indirect interactions will promote coexistence, however, facilitation may weaken the relationship between niche differences and complementarity if facilitated species persist at low abundances and contribute little to functioning (Fig. 1A); (4) Nitrogen addition and foliar pathogen removal will weaken the relationship between niche differences and complementarity, while strengthening the link between competitive ability differences and interspecific selection (dominance) effects (Fig. 1C). Our study extends links between coexistence mechanisms and biodiversity effects previously established in two-species communities to multispecies communities^8^. By taking this approach, we show that the conditions for stable coexistence are able to predict observed functioning across contrasting environmental contexts, providing a theoretical basis for predicting diversity effects under global change.

## Results and discussion

When facilitation was excluded (see methods), we found that communities with greater niche differences had higher complementarity effects and more negative intraspecific selection effects, while larger niche differences were associated with higher interspecific selection (dominance) effects (Figs. 2 and 3). Coexistence mechanisms also partially explained how fast-slow strategies, represented by specific leaf area (SLA), altered biodiversity effects. Communities dominated by slow-growing species showed greater niche differentiation and, consequently, higher complementarity and more negative intraspecific selection effects. However, nitrogen also increased competitive ability differences in these slow communities, which led to increased interspecific selection effects (Fig. 3). This highlights the importance of fast-slow functional strategies for understanding the causes and consequences of biodiversity change. However, traits also altered biodiversity effects in ways that could not be explained by the coexistence mechanisms (these are the direct effects of traits in structural equation model), although the strength and direction of these unexplained effects differed between nitrogen treatments and seasons. Importantly, including facilitation decoupled the relationship between coexistence mechanisms and biodiversity effects, showing that links between coexistence mechanisms and biodiversity effects depend on how species interactions were represented. Finally, although indirect interactions did not increase any of the diversity effects directly, niche differences calculated excluding indirect interactions (the pairwise feasibility domain) did not increase complementarity, suggesting that indirect interactions may contribute to this relationship. Below, we unpack these relationships step by step.

**Fig. 2.**
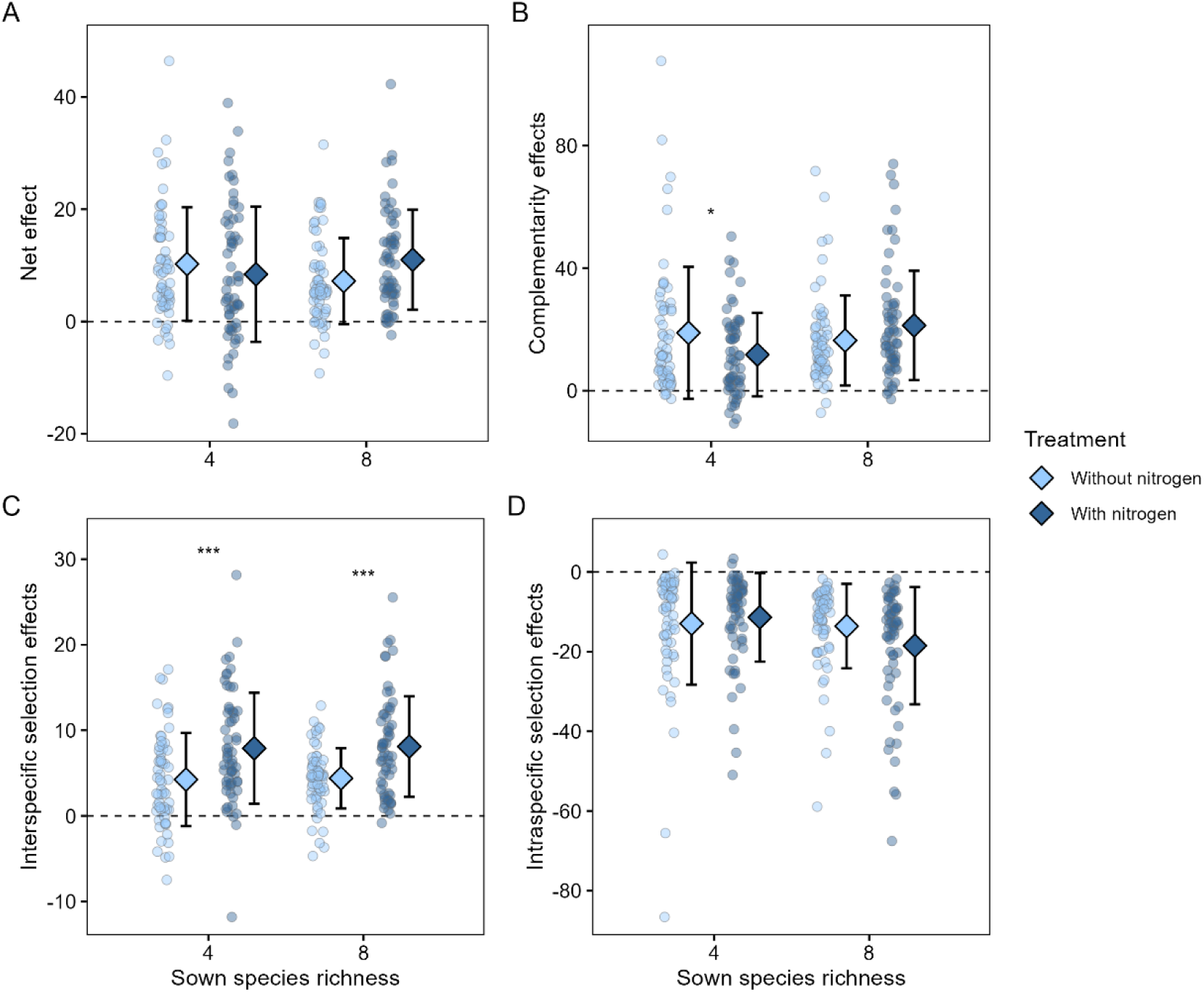
Mean diversity effects on productivity in four-and eight-species communities across seven years. Panels A-D show the mean net diversity effects, complementarity effects, inter-and intraspecific selection effects in four-and eight-species communities with and without nitrogen. Individual observations are shown alongside group means, with error bars representing ± 1 standard deviation (SD). Asterisks indicate differences between communities with and without nitrogen addition within each richness level, based on Tukey post-hoc comparisons (*, *p* < 0.05; ***, *p* < 0.001).

**Fig. 3.**
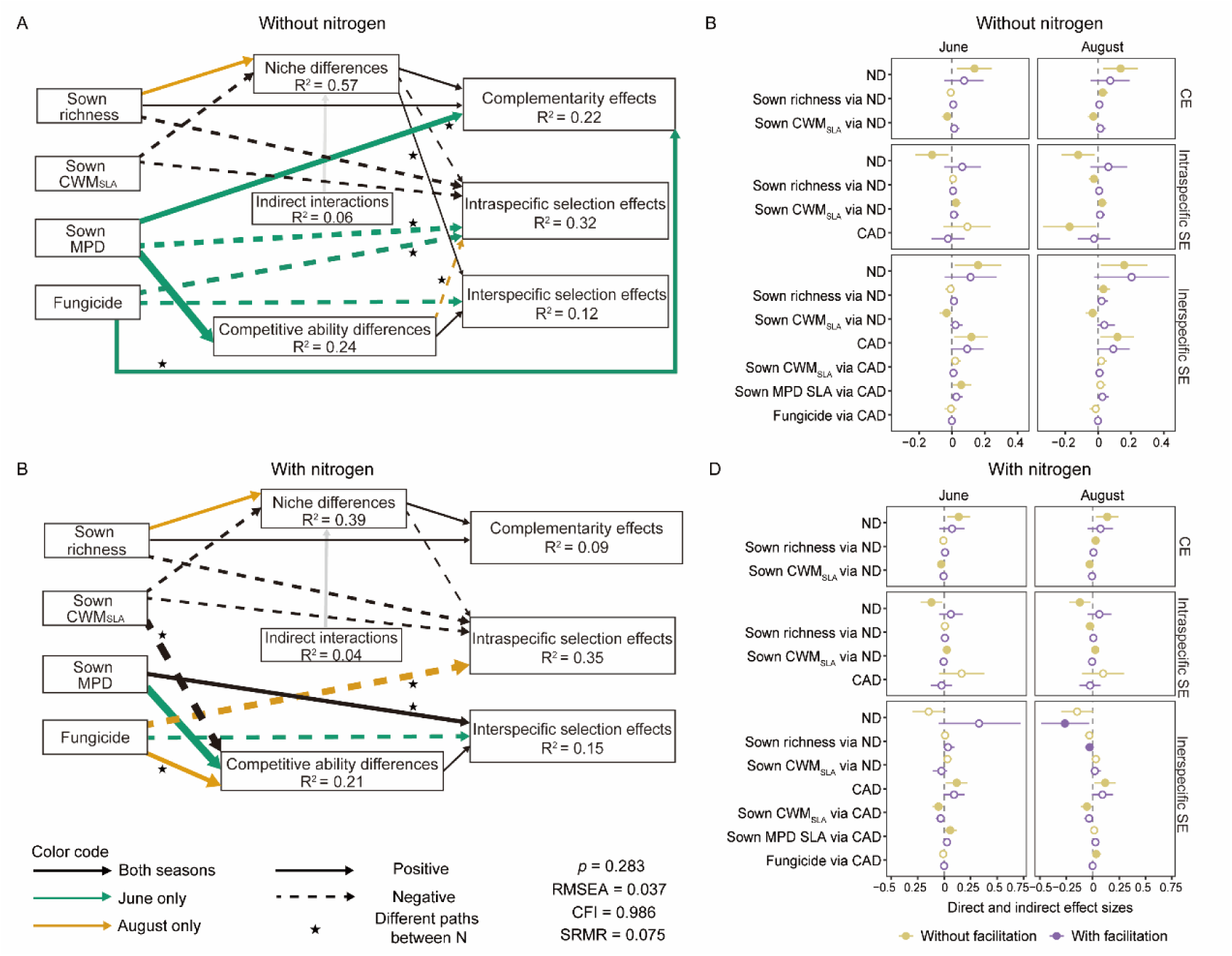
Relationships between coexistence mechanisms and biodiversity effects with and without facilitation. Structural equation model (SEM) linking sown species richness, functional composition (Sown CWM_SLA_), functional diversity (Sown MPD), fungicide, coexistence mechanisms (standardized niche differences, competitive ability differences, and indirect interactions) without facilitation based on sown species composition, and mean diversity effects (Complementarity effects, intraspecific and interspecific selection effects) without nitrogen (A) and with nitrogen (C). Solid and dashed arrows indicate significant positive and negative effects, respective (*p* < 0.05). Paths that are consistent across June and August are shown in black, season-specific paths are highlighted in bluish-green (June only) or golden-orange (August only). Although niche differences and indirect interactions are mathematically related in the structuralist approach, indirect interactions can theoretically promote, constrain or have no effect on coexistence^31^. Based on this theoretical rationale, the directional path from indirect interactions to niche differences is shown in grey. Arrow thickness is proportional to the strength of the path coefficient. Black stars indicate paths that differ significantly between nitrogen treatments. Non-significant and marginally significant paths, covariances, and path coefficients are omitted for clarity. R² values are shown for all endogenous variables, representing the maximum observed across two seasons without and with nitrogen. Model fit was adequate: CFI = 0.986, RMSEA = 0.037, SRMR = 0.075, and full model details are provided in Table S3. Selected direct and indirect effects from SEMs excluding or including facilitation without nitrogen (B) and with nitrogen (D) are shown. These include the direct effects of niche differences (ND) and competitive ability differences (CAD) on complementarity (CE), intraspecific selection effects (intraspecific SE), and interspecific selection effects (interspecific SE), and the indirect effects of sown richness, sown CWM_SLA_, sown MPD, and fungicide application mediated through ND or CAD. Points represent estimates, and error bars show 95% Monte Carlo confidence intervals.

## Diversity effects on productivity

We first calculated mean diversity effects on biomass for each plot, by averaging them across seven years, separately for early (June) and late (August) harvests. We analyzed the two harvests separately because variation in phenology can alter species interactions and contributions to biomass^46^. As in many studies, we found positive net diversity effects (mixtures produced more biomass than expected from the monocultures) (Fig. 2A), which were primarily driven by positive complementarity effects (Figs. 2B and S1), and, to a lesser extent, by positive interspecific selection (dominance) effects (Fig. 2C). Intraspecific selection effects were mostly negative (Fig. 2D) and therefore reduced net diversity effects, as expected from previous studies^20,55^. The negative intraspecific selection effects indicate low monoculture biomass species benefit disproportionately from growing in mixtures but without their increase in biomass affecting other species^52^.

We then determined the context dependency of biodiversity effects using linear mixed effects models to test how mean diversity effects depended on plant community characteristics: species richness, functional composition (community-weighted mean SLA based on the sown composition; Sown CWM_SLA_), functional diversity (mean pairwise distance in SLA among sown species; Sown MPD), nitrogen addition, foliar fungal pathogen suppression and season. Nitrogen strongly modified biodiversity effects and altered their response to species richness and functional diversity (Figs. 2 and S2, Table S1). Consistent with previous work^37,56,57^, nitrogen addition increased the dominance of high-biomass species (positive interspecific selection effects, Fig. 2C). Nitrogen also reduced complementarity in four-species communities and dampened the positive relationship between functional diversity and complementarity effects (Figs. 2B and S3). This is consistent with previous findings showing that nitrogen addition can weaken both complementarity effects and positive effects of functional diversity on multifunctionality^49,58^. Nitrogen also amplified the negative effect of species richness on intraspecific selection effects (Figs. 2D and S2). Strong intraspecific selection effects could be driven by less competitive species suffering strong negative frequency dependence, for instance, increased pathogen attack in monoculture, which might have been enhanced by nitrogen enrichment if nitrogen increased natural enemy attack^59^. The reduced intraspecific selection effects under fungicide application support this explanation (Fig. S2). We next tested whether coexistence mechanisms in the mixtures might explain variation in diversity effects between mixtures.

## H1: Coexistence mechanisms link to diversity effects when facilitation is removed

We tested whether functional diversity and composition influenced biodiversity effects observed on a plot, through their effects on coexistence mechanisms. For each plot, we calculated the structural niche difference, competitive ability difference, and the contribution of indirect interactions to coexistence, based on its sown composition (see methods). We used pairwise interaction coefficients and species intrinsic growth rates, measured under the same treatment combination (N x fungicide) and season in a phytometer experiment conducted at the same field site^53^. To compare coexistence mechanisms among plots with different species richness, we developed a randomization approach that reshuffled interaction coefficients and intrinsic growth rates between species (see methods). This allows us to calculate whether a given species composition had higher or lower niche differences, competitive ability differences and contribution of indirect interactions to coexistence, than expected under this randomization. Following previous studies^8,10,15^, we first calculated coexistence mechanisms without facilitation by setting positive interactions to close to zero (see methods), and later tested whether including facilitation altered links between coexistence mechanisms and biodiversity effects. As nitrogen and season modified the responses of biodiversity effects to functional diversity and composition, we fitted a multigroup structural equation model with four groups defined by nitrogen treatment and season (June without nitrogen, June with nitrogen, August without nitrogen and August with nitrogen) (initial full model and hypothesized causal relationships presented in Fig. S4). To assess whether relationships differed among groups, we compared models in which a given path coefficient was estimated separately for each group, with models in which that coefficient was constrained to the same value across groups (details in method)^60^. Here we focus on the relationships involving niche and competitive ability differences, the contribution of indirect interactions is discussed separately.

When facilitation was excluded, communities with greater niche differences had higher average complementarity effects (Figs. 3 and 4), consistent with theoretical expectations. At the same time, higher competitive ability differences led to greater interspecific selection effects, consistent with greater dominance by competitively superior species (Figs. 3 and 4). Greater niche differences were also associated with more negative intraspecific selection effects (Fig. 3), which agrees with the idea that niche differentiation disproportionally benefits low-biomass species in mixtures, without this coming at the expense of other species^52^. These links were consistent across nitrogen treatments and seasons (Fig. 3). In line with a recent study^15^, our findings therefore show that a statistical partitioning of biodiversity effects does in fact reflect underlying coexistence mechanisms in multispecies communities, regardless of environmental context. Our approach also provides a robust test of the link between coexistence mechanisms and biodiversity effects, because coexistence mechanisms estimated from interactions measured in a separate phytometer experiment can predict biodiversity effects in the main experiment. This use of independently measured interaction coefficients distinguishes our approach from previous experiments^8,15^.

**Fig. 4.**
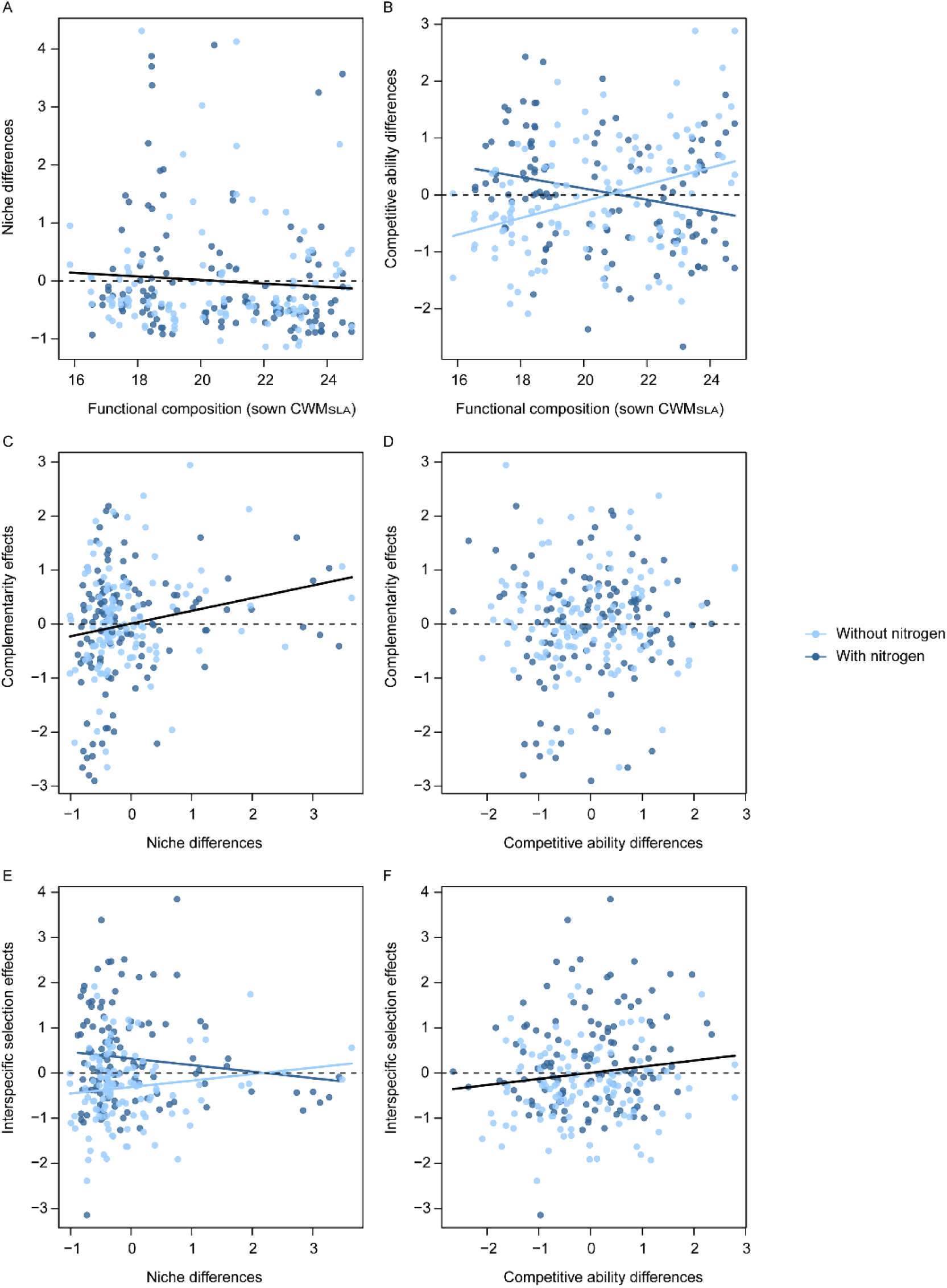
Relationships between (A) functional composition (sown CWM_SLA_) and niche differences, (B) functional composition (sown CWM_SLA_) and competitive ability differences, (C, E) niche differences based on the sown composition (without facilitation) and complementarity and selection effects, and (D, F) fitness differences based on the sown composition (without facilitation) and complementarity and selection effects. Graphs show partial residuals from the structural equation model. X-axes show unscaled functional diversity, functional composition, predicted niche differences, and competitive ability differences. Points represent partial residuals from SEM, and lines show the corresponding SEM-predicted relationships.

However, although communities with larger niche differences had greater complementarity effects, our models predicted that many of the sown species could not stably coexist. In detail, none of the full-community equilibrium solutions had positive abundances for every species based on interaction coefficients and intrinsic growth rates. These results are consistent with previous studies examining how coexistence mechanisms vary across environmental conditions^36,61,62^. This pattern arises because structural niche difference quantifies the size of the feasibility domain under the assumption of global stability, rather than local stability^31^. To further understand the lack of global stability for the communities we studied, we simulated population dynamics using a multispecies Ricker model. Although most of our communities are predicted to lose one species relatively fast (4-5 simulation steps when facilitation was excluded), we also observed that at least half of the sown species could persist for 200-time steps in 17.5% (June) and 34.2% (August) of plots (Fig. S5). Importantly, communities with larger niche differences were predicted to maintain more species (Table. S2). By relating these niche differences to diversity effects, our results show that competitive coefficients measured in a single year can explain statistically significant variation in the functioning of perennial grassland communities across multiple years. This finding suggests that the structure of species interactions may remain relatively consistent across time^63^. Although future work should explore the degree of temporal stability in plant-plant interactions, we hypothesize that any changes in species interactions were not sufficient to substantially alter competitive hierarchies or the balance between intra-and interspecific interactions across seven years in our perennial grassland. This might not be the case in systems with faster life cycles, such as annual plant communities^64^. Finally, our results suggest that population models, considering only changes in biomass, may be able to represent perennial plant community dynamics^65^.

## H2: Functional traits link to coexistence mechanisms and biodiversity effects

We also found that the coexistence mechanisms in our multispecies communities could, to some extent, explain how traits determined biodiversity effects (Figs. 3 and S6). In June, at peak biomass, communities containing a mix of slow-and fast-growing species (high sown MPD) had greater competitive ability differences and this led to higher interspecific selection effects (Fig. 3). This was consistent with the differences in intrinsic growth rates between fast-and slow-growing species in June. Functionally diverse communities also tended to have higher niche differences (*p* = 0.05, Fig. S6). Differences in SLA are expected to promote niche differences as fast-and slow-growing species use different strategies to acquire resources such as light and water^18,66,67^. However, our results suggest that these differences in resource use can also translate into differences in their overall competitive ability^66,68^. In addition to differences in traits, the functional composition of communities affected the type of interactions occurring and therefore the degree of coexistence (Fig. 3). Slow-growing communities (with low CWM_SLA_) had higher niche differences and this resulted in higher complementarity and more negative intraspecific selection effects (Fig. 3). By contrast, fast-growing species may be less differentiated in resource use, and this may have led to lower niche differences and lower complementarity (Fig. 3)^18^. A shift towards dominance by fast-growing species in intensively managed systems^16^, is therefore likely to reduce niche differences and the positive effects of diversity on functioning.

Several of the links between fast-slow traits and biodiversity effects were direct, meaning that they could not be explained by coexistence mechanisms estimated from pairwise interactions between the species (Fig. 3). In June, without nitrogen addition, communities with greater functional diversity (higher sown MPD) had higher complementarity effects and more negative intraspecific selection effects, independent of their differences in coexistence mechanisms (Fig. 3A). This result suggests that species with low monoculture biomass performed much better in functionally diverse mixtures. Fast-growing communities also had more negative intraspecific selection effects regardless of nitrogen addition (Fig. 3). As discussed above, one possible explanation is that some poorly defended fast-growing species suffer more from natural enemies in their monocultures but experience less damage and recover more in diverse communities. Coexistence mechanisms therefore only captured part of the trait effects, which is consistent with the idea that niche differences are necessary but not sufficient for complementarity effects^10^.

Complementarity effects, and the other diversity effects, also depend on the relative abundances of species in mixtures, and the direct trait effects suggest that there are additional mechanisms regulating species abundances that are not captured by pairwise competition coefficients. For instance, species may be able to coexist stably by being strongly niche differentiated from others but might only occur at low abundance in communities if they suffer from very strong negative frequency dependence caused by specialist pathogens^69–71^. Another mechanism regulating abundances could be higher-order interactions that occur when the presence of a third species modifies the interaction between a pair of species^72^. These differ from indirect interactions and cannot be readily incorporated with the structural approach we used^31^. Finally, although SLA captures an important axis of plant functional variation in our study system^20,48,73^, other traits, particularly those more directly linked to underlying mechanisms of species interactions, such as resource uptake or interactions with specialist microbes, may further improve prediction of community dynamics, coexistence mechanisms and their consequences for ecosystem functioning^74^. Together, our results suggest that fast-slow functional strategies influence diversity effects not only via changes in coexistence mechanisms, but also through additional mechanisms not captured by pairwise competition coefficients assumed to be time-invariant.

## H3: Influence of facilitation and indirect interactions on links between coexistence mechanisms and biodiversity effects

Facilitation can promote species coexistence and may also generate positive complementarity effects^26,75,76^, however, studies linking coexistence and functioning have rarely considered facilitation explicitly. Thanks to the theoretical framework used here, we can test how facilitation altered relationships between coexistence mechanisms and biodiversity effects. To include facilitation, we recalculated coexistence mechanisms for each plot using the full range of interaction coefficients^53^, rather than assigning positive interactions close to zero, as has been done in many previous studies. When facilitation was included, the positive effect of niche differences on complementarity effects and the negative effect of niche differences on intraspecific selection effects were no longer supported (Figs. 3, S6 and S7; Table S3). Even the positive relationship between competitive ability differences and interspecific selection effects was weakened (Figs. 3, S6 and S7). Therefore, the relationships between coexistence mechanisms and biodiversity effects vanished in our study once facilitation was included.

Including facilitation increased both the observed and standardized niche differences, particularly in fast-growing communities (Figs. S8 and S9). However, the niche differences including facilitation failed to predict the persistence of most of the sown species in the simulated population dynamics using a multispecies Ricker model (Fig. S5 and Table S2). This lack of stability when facilitation was included suggests that our approach did not capture additional mechanisms that are important for understanding how interactions shift from facilitation to competition across years, life stages, and population densities^77^. For instance, facilitation may occur in particular years or benefit juveniles at low population densities, but shift towards competition as environmental conditions improve, individuals reach adulthood, and population densities increase. Without accounting for these context-dependent shifts, including facilitation could lead to unrealistic predictions in which facilitated species eventually become dominant and exclude other species^76^. Further, in our experiment, facilitation was observed mainly among the fast-growing species^53^, perhaps because they benefited more from microclimate amelioration. However, fast-growing species are generally less competitive in our site^53^ and the species that benefited most from facilitation had the lowest relative abundances in mixtures and contributed least to complementarity (e.g., *Rumex acetosa*) (Figs. 5 and S10). This result aligns with earlier findings from the Jena biodiversity experiment, where less abundant species also did not show stronger overyielding in diverse plots^78^. Our results suggest that although facilitation theoretically promotes coexistence by buffering low abundance species from extinction, it does not necessarily translate into higher complementarity because facilitated species that remain rare in mixtures contribute little to complementarity effects.

**Fig. 5.**
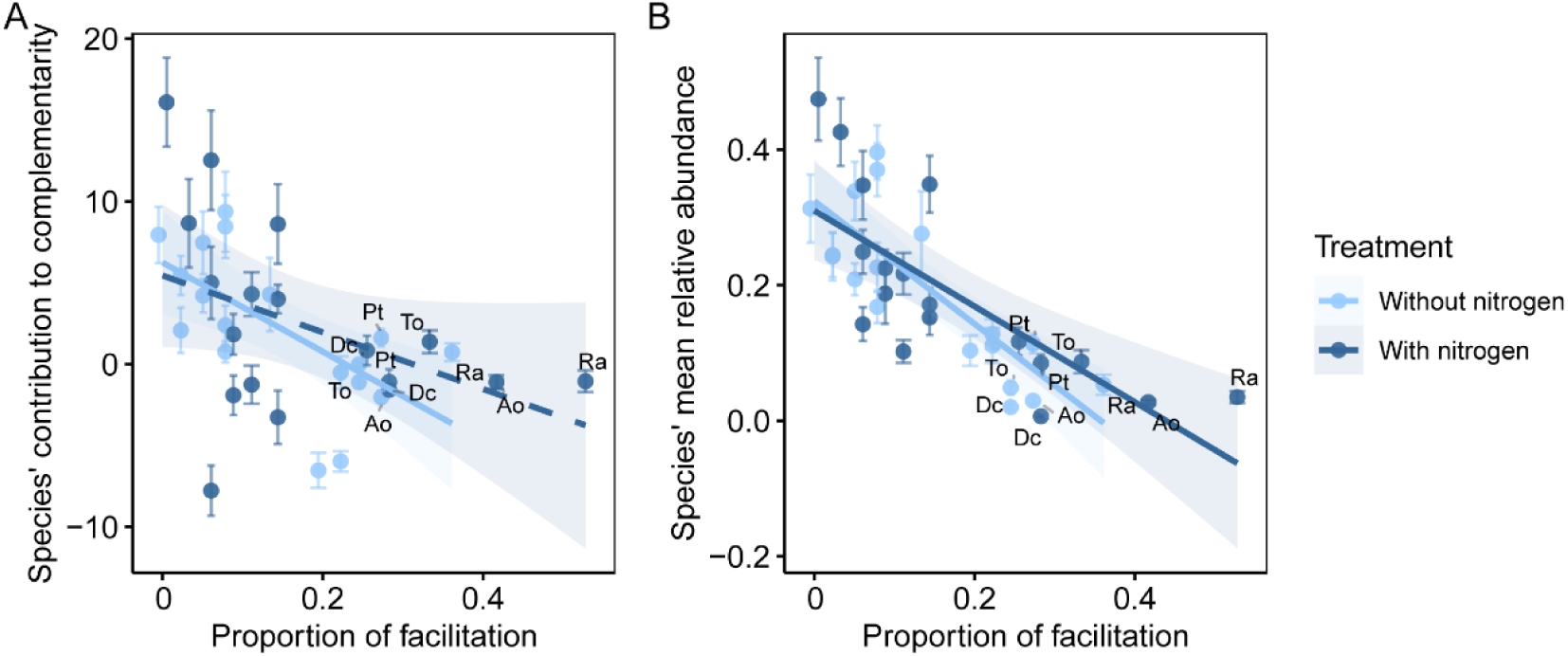
Relationships between species’ contributions to complementarity effects (ΔRYᵢ × Mᵢ) and the proportion of facilitation each species received (A), and between species’ mean relative abundance in mixtures and the proportion of facilitation received (B) in June. Solid lines represent significant (*p* < 0. 05) fit from linear regressions across treatments with shaded areas represent 95% confidence intervals. To improve readability, only the top five species receiving the highest levels of facilitation are labeled. Species abbreviations are listed here: Ra: *Rumex acetosa*; Ao: *Athoxanthum odoratum*; Dc: *Daucus carota*; To: *Taraxacum officinale*; Pt: *Poa trivialis*. In (A), species that received more facilitation had lower complementarity effects (With nitrogen: *p =* 0.08, R^2^ = 0.13; Without nitrogen: *p <* 0.01, R^2^ = 0.38). In (B), species that received more facilitation had lower relative abundances (With nitrogen: *p* < 0.001, R^2^ = 0.54; Without nitrogen: *p* < 0.001, R^2^ = 0.65).

We were also able to test the effect of indirect interactions on diversity effects^24,25^. In the structuralist approach, the difference between the size of the full-community feasibility domain and that of the domain allowing all species pairs to coexist (the so-called community-pair differential, D) measures the contribution of indirect interactions to coexistence^31^. The observed differential was often positive, particularly in four-species communities (Fig. S11), which means that indirect interactions contribute to coexistence. We next tested whether the increase in niche differences due to indirect interactions led to higher complementarity by calculating niche differences without indirect interactions, i.e., replacing the standardized structural niche differences in the structural equation model with the feasibility domain allowing all species pairs to coexist (standardized pairwise feasibility, which explicitly excludes the effect of indirect interactions). The pairwise niche differences (without indirect interactions) still led to more negative intraspecific selection effects but had no effect on complementarity effects (Fig. S12 and Table S4). This suggests that indirect interactions did increase complementarity by indirectly increasing niche differences. This effect seemed to be especially strong in eight species communities, where the standardized pairwise feasibility was near zero and where indirect interactions strongly increased multispecies niche differences (Fig. S13). Indirect interactions therefore seem to be important for coexistence and complementarity in more diverse communities. Recent mesocosm experiments have found positive effects of indirect interactions on productivity and ecosystem multifunctionality^79,80^. Our results suggest that although indirect interactions did not increase biodiversity effects directly, they may indirectly promote the positive effect of niche differences on complementarity effects in multispecies communities.

## H4: Effects of nitrogen, pathogens and seasons on coexistence mechanisms and biodiversity effects

Nitrogen altered links between traits, coexistence mechanisms, and biodiversity effects in three ways. First, nitrogen modified the effect of fast-slow strategies on coexistence mechanisms. Without nitrogen addition, competitive asymmetries tended to be greater among fast-growing species (*p* = 0.08, Fig. S6). However, nitrogen reversed this relationship and resulted in greater competitive asymmetries among slow-growing species (Fig. S6). One possible explanation is that nitrogen induced limitation by other resources (e.g., water and phosphorus)^81,82^, favoring some slow-growing species with deeper rooting depths. The marginally significant positive relationship between functional diversity and niche differences remained stable across nitrogen treatments (*p* = 0.05, Fig. S6), in contrast to our expectation that nitrogen would weaken the relationship by reducing niche dimensions and intensifying light competition^83–85^ (Fig. 3). This may imply that fast-slow strategies promote niche differentiation through other underlying mechanisms such as different responses to insect herbivores, soil microbes or other limiting nutrients.

Second, nitrogen altered two links between coexistence mechanisms and biodiversity effects. Higher niche differences tended to reduce interspecific selection (dominance) effects under nitrogen addition but increased interspecific selection effects without N (Fig. S6). This is because nitrogen increased interspecific selection effects mainly in communities with low niche differences (Fig. 4G). This agrees with the idea that nitrogen enrichment amplifies competitive asymmetry particularly when species overlap strongly in resource use (when niche differences are small), leading to stronger dominance effects^53^. By contrast, greater stabilizing niche differences may buffer inferior species and constrain the extent to which high-performing species dominate communities without nitrogen addition, which explains why we observed a positive association between niche differences and dominance effects^8^. In August, during the regrowth after mowing, higher competitive ability differences led to more negative intraspecific selection effects without nitrogen addition but not with nitrogen addition (Fig. 3). The dampening of both effects may occur because nitrogen generally increases competitive dominance and allows high biomass species to dominate.

Finally, the direct effect of functional traits and fungicide application on diversity effects varied with nitrogen treatment and season. In June, at peak biomass, complementarity effects were higher and intraspecific selection effects were more negative in functionally diverse communities and when foliar fungal pathogens had been reduced by fungicide application (Fig. 3A). Nitrogen addition weakened these effects (Fig. 3B). As we discussed earlier, nitrogen enrichment may strengthen associations between fungal pathogens and their host plants, reducing the extent to which susceptible fast-growing species benefit from growing in diverse mixtures^50^. Fungicide application also reduced interspecific selection effects in June under both nitrogen treatments. In August, during post-mowing regrowth, fungicide application had two distinct effects in nitrogen-enriched plots. Fungicide application had a direct negative effect on intraspecific selection effects, with more negative values in plots received fungicide application. It also increased interspecific selection effects by increasing competitive ability differences (Fig. 3B). However, fully understanding these seasonal specific effects would require more information on the low-level processes driving coexistence and diversity effects^9^. These three sets of modifications highlight that nitrogen alters not only the links between functional strategies and coexistence mechanisms but also direct trait effects, which then changes biodiversity effects. This calls for more studies on the context dependency of these links, if we are to predict changes in functioning using traits and coexistence mechanisms.

Taken together, our study provides empirical evidence on the extent to which diversity effects can be predicted by underlying species coexistence mechanisms. With two large independent experimental studies and complex statistical modeling, we were able to link functional traits and coexistence mechanisms to biodiversity effects and demonstrate that coexistence mechanisms estimated from local species interactions in one year did predict diversity effects on plots across time. This hard test addresses the concern that coexistence models are rarely externally validated, by showing that interactions measured in one context can be used to predict biomass in another context^86^. Further, differences in interactions between species differing in fast-slow traits led to different coexistence mechanisms and diversity effects depending on fast-slow functional diversity and composition. However, these relationships broke down when we included facilitative interactions because facilitation increased niche differences but not complementarity. We also found that some effects of fast-slow traits on diversity effects were not explained by coexistence mechanisms. Both these results may occur because coexistence mechanisms predict how many species can persist in communities, but they do not entirely predict species’ abundances and evenness, therefore they cannot explain all the variation in diversity effects between communities. Although the overall links between coexistence and biodiversity effects were stable across nitrogen treatments and seasons, many effects of traits and pathogens differed, suggesting that species’ abundance distributions are particularly variable across contexts. Understanding biodiversity effects therefore requires considering not only the interactions that shape coexistence but also the processes that influence species abundances under changing resource and environmental conditions. However, our results point the way to a framework uniting coexistence theory and biodiversity effects in communities undergoing global change.

## Materials and methods

### The PaNDiv experiment

The PaNDiv experiment was established in October 2015 in Münchenbuchsee (Bern, Switzerland). It consists of 216 2 × 2 m plots in which different communities were established, varying in species richness (1, 4, 8 and 20 species), functional composition and functional diversity, nitrogen enrichment (in the form of urea; 0, 100 kg N per ha per year) and foliar fungicide treatment (Score Profi by Syngenta Agro AG, 24.8 % difenoconazole and Ortiva by Syngenta Agro GmbH, 22.8 % azoxystrobin). The 20 species plots were not included in this analysis, see reasoning below. Plots were separated by 1m grass paths and mown twice a year, reflecting typical extensive grassland management in this region. Twenty perennial grassland species were used in the experiment (Table S5), ranging from fast-growing to slow-growing species based on their specific leaf area (SLA) and leaf nitrogen content. Species combinations were randomly selected from the respective species pool (i.e. fast, slow or all). Each combination received the four treatments (fungicide × nitrogen), randomly allocated within blocks (4 in total). Plots were weeded three times annually (April, July and September) to maintain the intended species compositions. A detailed description of the experimental design is available in earlier studies^48,50^.

### Productivity, trait composition and diversity

The percentage cover of all sown species, bare ground and weeds was visually estimated twice a year (June and August) in all plots. After each cover estimate, aboveground biomass was harvested from two 50 × 20 cm subplots within the 1 m^2^ core area. Biomass per species was estimated by multiplying total plot biomass (weeds were excluded) by each species’ relative cover. A prior comparison showed that species estimated biomass (total biomass without weeds × relative cover of target species) was highly correlated with the sorted biomass the (R^2^ = 0.994)^49^. On average, weed cover accounted for around 5% of total cover, therefore removing weeds had minimal effect on biomass estimates^20,48^. Biomass was collected using this method in June and August from 2017 to 2023, resulting in 14 sampling events. SLA was measured from 2016 to 2020 following a standardized protocol^87^ by collecting one leaf from 5 individuals in each monoculture plot across four treatments. SLA per species was calculated by averaging five samples and five years. Community weighted mean SLA (hereafter, Sown CWM_SLA_) was calculated per plot using sown relative abundance: 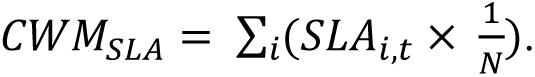 Where *N* is sown richness and SLA*_i_*, *_t_* is the SLA of species *i* in treatment *t*. Sown CWM_SLA_ indicates the fast-slow growing strategy of a community. Functional diversity was calculated as the mean pairwise distance in SLA (hereafter, Sown MPD) following an earlier study^88^.

### The phytometer experiment

To estimate intra-and interspecific competition and intrinsic growth rates, a phytometer experiment was conducted in 2019. Seedlings were transplanted into neighborhoods varying in density (from no neighbors to crowded) and species composition (intra-or interspecific neighbors) under four treatments. Each neighborhood was defined as a 20 cm radius around the focal phytometer plant^36^. Seedlings were germinated with commercial seeds (UFA Samen in Switzerland and Rieger Hoffmann in Germany) in a greenhouse in Ostermundigen (Bern, Switzerland) in August 2019. Two species, *Anthriscus sylvestris* (As) and *Heracleum sphondylium* (Hs), had poor germination rates and were excluded from the experiment. However, these two species also did not establish well on the PaNDiv experiment^20^. The remaining 18 species were planted into 10 different intraspecific neighborhoods at varying densities in the corresponding monoculture plots (intraspecific competition), three interspecific neighborhoods (interspecific competition), and empty neighborhoods (no competition) in the PaNDiv experiment. Phytometer biomass was harvested twice in June and August 2020). In total, 4’248 phytometers were planted; 91.5% (3’888) and 64.1% (2’722) were successfully recovered and harvested in June and August 2020, respectively. Interaction coefficients were estimated separately for each focal species using negative-binomial models, and therefore population dynamics were modelled with a Ricker model to have full correspondence between parameter estimation and population dynamics. Phytometer dry biomass was modelled as a function of the cover of each of the 18 neighboring species, treatment and season, with weed cover as a covariate. This approach therefore estimated the effect of each neighboring species while accounting for the other species present in the neighborhood. It allowed us to quantify both competition and facilitation. Following this, all intraspecific interaction coefficients (α_ii_) and pairwise interspecific interaction coefficients (α_ij_) were estimated per treatment and sampling season (June and August, respectively), i.e. 8 separate 18 × 18 interaction matrices. The effects of treatments (N × Fungicide), functional traits and season on competition networks are presented in the original paper^53^. We also measured intrinsic growth rates (λ) by planting phytometers of all 18 species in extra 2 × 2 m plots (16 in total) located next to the PaNDiv experiment without competition (competitors were suppressed using woven landscape fabric covering the bara ground). In each extra plot, we planted one phytometer of each species 20 cm apart. The 16 extra plots received the same four N × Fungicide treatments (control, nitrogen addition, fungicide application and the combined treatment of nitrogen and fungicide) and were maintained in the same way as the PaNDiv experiment.

## Data analysis

### Tripartite partitioning of diversity effects on productivity

To decompose diversity effects on aboveground biomass, we employed Fox’s tripartite partitioning framework^52^, which partitioned the net diversity effect (the difference between observed and expected total yield, ΔY) into complementarity effects, interspecific selection effects, and intraspecific selection effects. These correspond to Fox’s trait-independent complementarity, dominance effects and trait-dependent complementarity, respectively^52^. This framework retains the complementarity component of Loreau and Hector’s additive partitioning while further separating the selection effect into interspecific (dominance) and intraspecific selection (trait-dependent complementarity) effects ^51^. Net diversity effect can be expressed as:

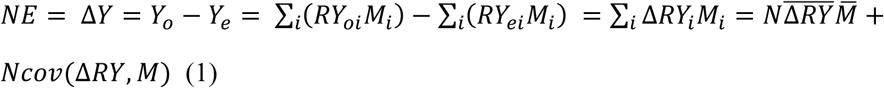

Where RY_oi_ is the observed relative yield of species *i* in mixture (equals to *Y_i_*⁄*M_i_*), and *M_i_* and *Y_i_* a are yields of species *i* in monoculture and mixture, respectively. *RY_ei_* is the expected relative yield of species *i* in mixture, which equals the sown relative abundance (1/*N*, N is sown richness). Δ*RY_i_* is the difference between observed and expected relative yields of species *i*. Symbols *X̄*and *cov*(*X*, *Y*) represent the mean and covariance of corresponding variables, respectively. The first term on the left-hand side of equation (1) quantifies complementarity effects, while the second term quantifies selection effects. In the tripartite partitioning framework, the “selection effect” term was further decomposed:

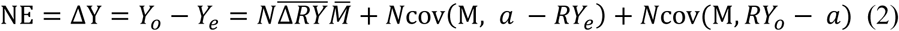

Here, *a_i_* is the observed relative cover of species *i*, used as a measure of its relative abundance, *RY_oi_* = *Y_oi_* ⁄ *M_i_* is its observed relative yield, and *RY_ei_* = 1/*N* is its expected relative yield. In equation (2), the first term equals the “complementarity effect” in Loreau and Hector^51^. It quantifies the average deviation of observed relative yields from their expected values, multiplied by species richness and mean monoculture yield. The first covariance term is the covariance between species’ monoculture yields and the differences between observed and expected frequencies (relative abundances), with a positive value indicating high monoculture yield species dominate at the expense of species with low monoculture yields (dominance effects sensu Fox). The second covariance term is the covariance between monoculture yield and the deviation of observed relative yield from observed frequency. Following Cappelli et al.^20^, we refer to these as interspecific and intraspecific selection effects to make explicit that both are components of the original selection effects. The interspecific term indicates effects arising from changes in one species frequency that come at the expense of other species and the intraspecific term indicates that changes in the biomass of one species in mixture do not affect the other species. Intraspecific selection effects are negative when species with low monoculture yields attain high observed relative yields and/or species with high monoculture biomass have lower relative yields than expected from their frequency in the mixture, i.e., not at the expense of other species. The sum of the two covariance terms is the “selection effect” in Loreau and Hector^51^. Because species biomass was estimated from relative cover and total plot biomass, our approach does not independently measure species-specific variation in biomass production per unit cover.

We calculated diversity effects for 4 and 8 species plots at each sampling point (year and season) using the mean aboveground biomass per sampling subplot (50 × 20 cm). As we described above, *A. sylvestris* (As) and *H. sphondylium* (Hs) were excluded from the analysis due to their poor establishment. *Daucus carota* (Dc) had extremely low monoculture biomass across all four treatments (N◊F) in June 2017, leading to extremely high complementarity effects and low intraspecific selection effects. We therefore excluded *D. carota* from the calculation of diversity effects for this harvest and recalculated the number of species included in each community. We subsequently removed extreme positive complementarity effects and negative intraspecific selection effects for three additional-harvest observations (three plots), for similar reasons. These observations were excluded as outliers because their partitioned components of biodiversity effects were sensitivity to low monoculture biomass values. However, no plots were excluded as we had information on diversity effects in those plots from other years. Mean diversity effects (Net effect, complementarity effects, interspecific and intraspecific selection effects) were calculated per plot and season, by averaging values across the seven years.

### Coexistence in multispecies communities

We used the structuralist approach to quantify the ability of the communities to maintain diversity according to two metrics: structural niche differences (Ω) and competitive ability differences (θ)^31^. In this approach, Ω reflects the so-called feasibility domain, and it is calculated via the interaction matrix including intra-and interspecific coefficients obtained from the phytometer experiment. Specifically, the feasibility domain indicates the range of intrinsic growth rates compatible with persistence (positive abundances) of all species within the community. Conversely, θ measures the angular deviation of the intrinsic growth rates from the feasibility domain’s centroid, and it can therefore be seen as a structural analogue of competitive ability differences between species. The centroid represents a reference growth rate vector with the greatest tolerance to small random perturbations while maintaining feasibility. Coexistence occurs when Ω exceeds θ.

This approach also allows us to quantify the contribution of indirect interactions to coexistence in multispecies communities. Specifically, the metric is called community-pair differential (D, indicated as Δ in the original paper^31^), measuring the degree to which coexistence is more or less easily obtained with the inclusion of indirect interactions. D is calculated as the difference between the size of the feasibility domain for the full community (Ω) and the size of the intersection of all pairwise feasibility domains (Ω_all). Ω_all measures the range of intrinsic growth rate combinations for which every species pair is feasible when considered in isolation. It is important to note that D ranges from-1 to 1, and more positive values indicate that indirect interactions provide more opportunities of coexistence^31^.

We computed Ω, Ω_all, θ, and D for each 4 and 8 species community based on the sown species composition of the plot. We excluded the 20 species plots because they all have the same sown species composition, and therefore we have no variation in coexistence mechanisms between 20 species plots with the same treatment. In each case we used the interaction matrix for the sown species from the corresponding N ◊ F treatment and season. In our study, although competition was prevalent, facilitation was observed for some fast-growing species^53^. We therefore assessed coexistence mechanisms both with and without facilitation to test whether facilitation changes the strength of links between coexistence and diversity effects. To exclude facilitation, we replaced positive interactions with randomly generated negative values that were close to 0 (< 10^-3^).

Because structural coexistence metrics inherently vary with species richness^36^, they cannot be directly compared across communities with varying richness. To enable comparison of coexistence mechanisms across all 4 and 8 species communities, we developed a null model to estimate expected Ω, Ω_all, θ and D for each community based on its sown composition. This model randomly shuffled species identities in the matrices of intrinsic growth rates and competition coefficients, under a given treatment and season. We ensured species identities remained consistent between the two matrices, which preserved the inherent connection between competition coefficients and intrinsic growth rates. We conducted 99 randomizations per community, recalculating Ω, Ω_all, θ and D based on randomized metrics. We then calculated standardized Ω, Ω_all, θ and D per plot, using mean and standard deviation of expected values:

Standardized Ω = (Observed Ω – Mean expected Ω) / SD of expected Ω

Standardized Ω_all = (Observed Ω_all – Mean expected Ω_all) / SD of expected Ω_all

Standardized θ = (Observed θ – Mean expected θ) / SD of expected θ

Standardized D = (Observed D – Mean expected D) / SD of expected D

This allowed us to calculate whether observed coexistence mechanisms were higher or lower than expected for that number of species and therefore to compare values across communities differing in species number. To test whether species persist longer in communities with greater niche differences, we additionally used a multispecies Ricker model to predict abundances of sown species in each plot for up to 200-time steps. The Ricker model has been extensively used to assess population dynamics and species coexistence under a community context^61,89^. For each plot and season, we ran simulations with and without facilitation using competition coefficients and intrinsic growth rates estimated per treatment (N × F) and sampling season. Total initial abundance was set to 1 per plot and was divided equally among the *N* sown species included in the simulation, giving each species an initial abundance of *1 / N*. We defined species loss as the first time a species simulated abundance fell below 0.1% of its initial abundance and were subsequently held at zero. We recorded the first step at which any species crossed the extinction threshold and the first step by which at least half of sown species had crossed it. If a simulation completed all 200 steps without reaching a given endpoint (persistence of all sown species), we assigned that endpoint a time of 200 steps. We then fitted linear mixed-effect models for the steps of the first species lost and of at least half of the sown species lost with and without facilitation. The time steps were log-transformed. We included standardized niche differences and seasons as the fixed effects, with block, sown species composition and plot as random effects.

### Linking coexistence to diversity effects

We first used linear mixed-effect models (LMMs) to assess how mean diversity effects (net diversity effect, complementarity effects, interspecific selection and intraspecific selection effects) were related to sown species richness, functional composition (sown CWL_SLA_) and diversity (sown MPD), nitrogen addition and pathogen removal and how they changed across two seasons. We included plot number, block and species composition as random effects. Complementarity effects and intraspecific selection effects were transformed using signed square-root transformation to improve residual normality and homogeneity of variance. Model assumptions were assessed using residual Q-Q plots and plots of residuals against fitted values (Fig. S14). All LMMs were conducted in R using the “lme4” package^90^.

To examine how coexistence mechanisms (the standardized Ω, θ and indirect interactions) link to biodiversity effects across seasons and nitrogen treatments, we fitted multigroup structural equation models with four groups defined by nitrogen addition and season (June without nitrogen, June with nitrogen, August without nitrogen and August with nitrogen) with the “lavaan” package in R^91^. Following the hypothesized causal relationships in Figure S4, the models included three sets of predictors: (1) sown richness, sown CWM_SLA_ and MPD; (2) fungicide application (3) standardized Ω, θ and indirect interactions, based on sown species composition. To assess whether relationships are consistent across N treatments and seasons, we compared the model in which each coefficient was estimated separately across groups with models in which it was constrained to be equal across groups, using a likelihood-ratio chi-square test. We ranked paths by the p-values from these initial single-path comparisons, from largest to smallest, and considered constraints sequentially in that order. At each step, we retained a constraint if it did not significantly reduce model fit; otherwise, the path remained unconstrained at that step. We first tested equality constraints across all four groups in a stepwise manner. For paths that could not be constrained across all groups, we then tested whether coefficients could be constrained between June and August regardless of nitrogen treatments, or between nitrogen treatments regardless of seasons. The final model retained all equality constraints accepted during this sequential procedure, while allowing the remaining coefficients to vary among groups. We quantified indirect effects of sown species richness, functional composition (sown CWM_SLA_) and diversity (sown MPD) and fungicide application on biodiversity effects through each coexistence mechanism as the product of the corresponding path coefficients in the final multigroup structural equation model. We estimated 95% confidence intervals using 10,000 Monte Carlo draws from the joint sampling distribution of the estimated coefficients, while retaining estimated covariances, with the “manymore” package in R^92^.

Although niche differences and indirect interactions are mathematically coupled in the structuralist approach, indirect interactions can theoretically promote, constrain or have no effect on coexistence^31^. Based on this theoretical rationale, we retained the directional path from the contribution of indirect interactions to niche differences in our SEM. However, we could also argue that this relationship is potentially spurious as the contribution of indirect interactions is calculated using the structural niche difference. Therefore, to better isolate the effect of indirect interactions on diversity effects, we also replaced standardized niche differences with standardized pairwise niche differences to assess whether relationships with diversity effects depend on using the full-community feasibility (standard niche differences) or the conditions permitting coexistence of all species pairs without accounting for indirect interactions (standardized pairwise niche differences). At the species level, we quantified the extent to which each species benefitted from facilitation (the proportion of interspecific facilitation) and related to species’ mean relative abundance in mixtures and its contribution to complementarity effects. We fitted separate linear regressions for each season with and without nitrogen addition. All analyses were performed in R 4.4.1^93^.

## Author contributions

N.A.P., S.L.C. and E.A. designed and set up the PaNDiv Experiment. E.A. and O.G. conceptualized the project. X.L., C.D., O.G. and E.A. contributed to the methodology. C.D., S.L.C. and N.A.P. collected the data. X.L. analyzed the data and wrote the first manuscript with substantial input from E.A. and O.G. All authors contributed to the review and revision of the manuscript.

## Data availability

All data and code that support the findings of this study are available in a public GitHub repository at this address: https://github.com/XiangyuLiu2011/Linking_coexistence_mechanisms_and_biodiversity_effe cts_in_multispecies_communities

## Acknowledgments

We are grateful to the whole PaNDiv team, especially Sylvain Chartier, Mervi Laitinen, Hugo Vincent, and all the helpers who contributed to the maintenance and data collection of both the phytometer and biodiversity-functioning experiments. We thank two anonymous reviewers for their insightful comments and suggestions. PaNDiv was funded by the Swiss National Science Foundation (Awards 31003A_160212, 310030_185260 and 310030_215442).

## Notes

### Competing Interest Statement

The authors have declared no competing interest.

