## Supplementary file for "Linking species coexistence mechanisms to biodiversity effects on productivity in multispecies communities under environmental variation"

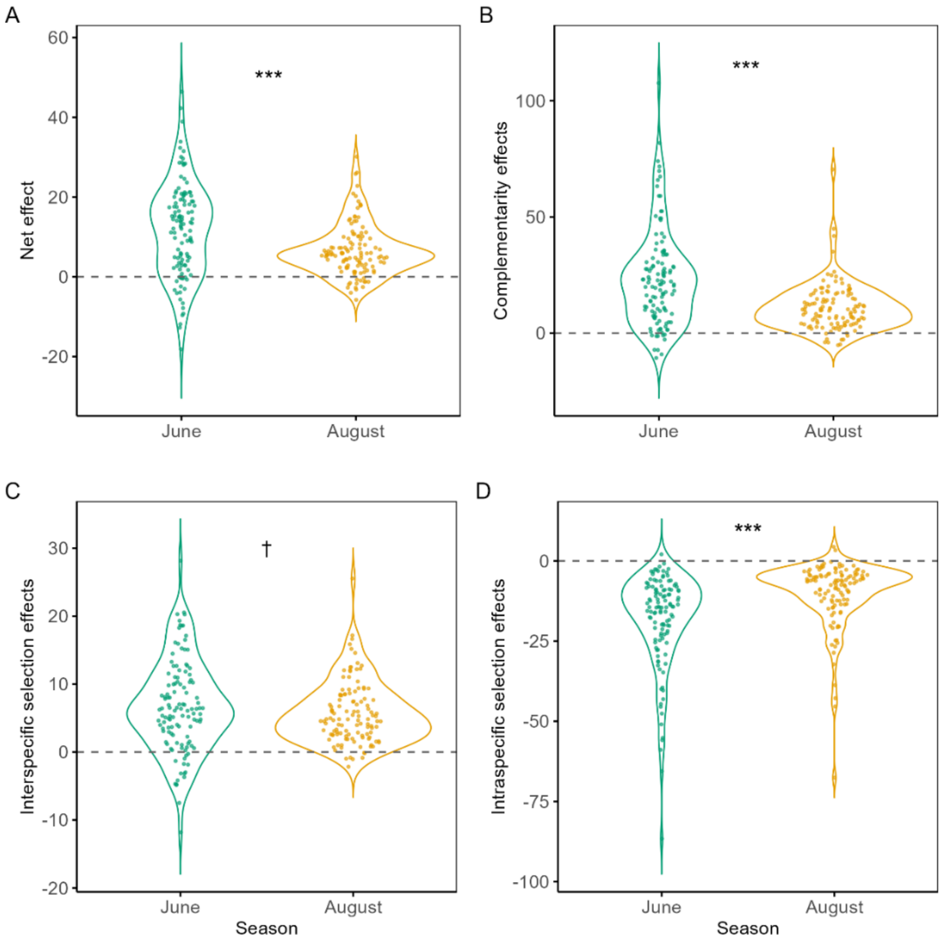


**Fig S1.** Mean biodiversity effects on biomass across seven years, separately for June and August. Panels A-D show the mean net diversity effects, complementarity effects, and inter- and intraspecific selection effects in June and August, respectively (raw data shown). Asterisks (*, $p$ < 0.05; ***, $p$ < 0.001) indicate significant differences between June and August based on Tukey post-hoc comparisons, marginally significant differences (0.05 ⩽ $p$ < 0.10) are marked with a dagger (†).

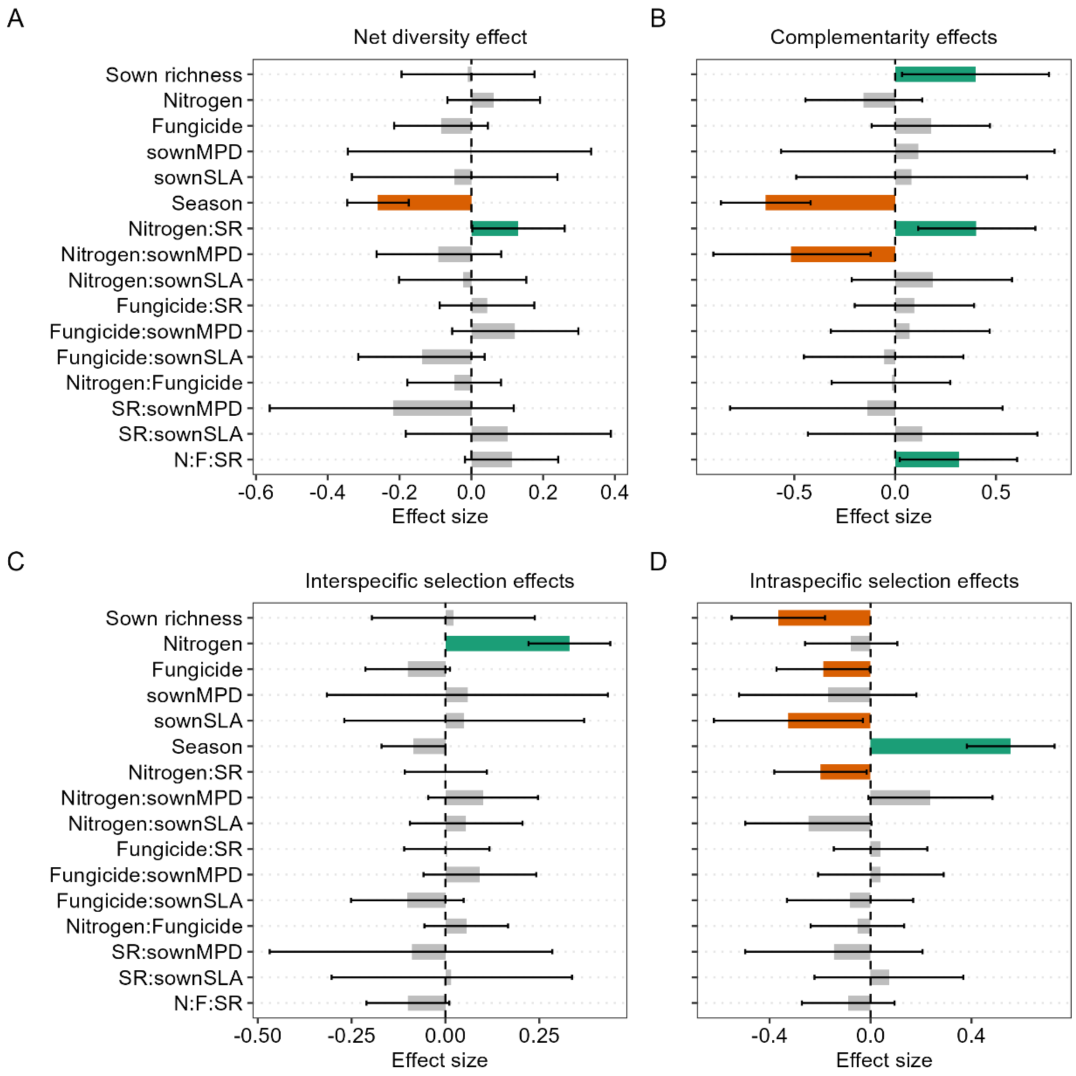


**Fig S2.** Results of linear mixed effect models showing drivers of mean diversity effects: (A) net diversity effects, (B) complementarity effects, (C) interspecific selection effect and (D) intraspecific selection effects. Shown are mean standardized effects (± confidence intervals) of each scaled and centered variable on diversity effects. Positive effects are shown in green, negative effects in orange, and non-significant effects are in gray. SR: sown species richness, Sown MPD: mean pairwise distance in SLA based on sown composition, Sown $CWM_{SLA}$: community weighted mean of SLA based on sown composition, N: nitrogen, F: fungicide.

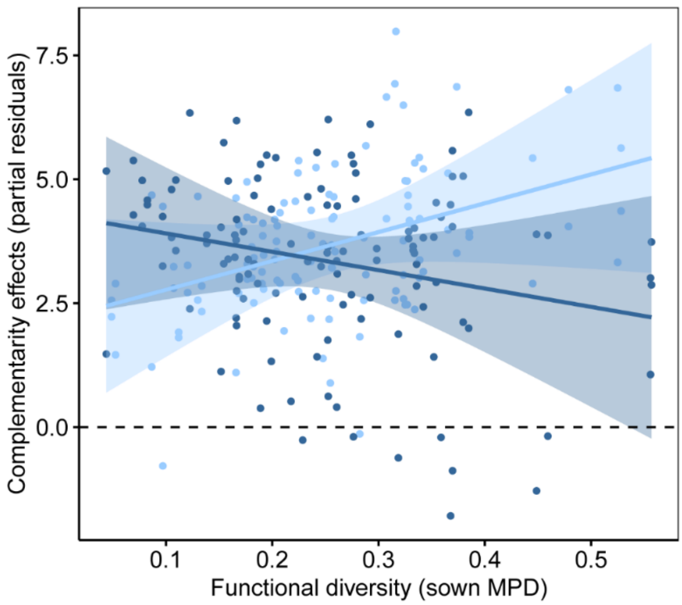


**Fig S3.** The partial effects of sown functional diversity (Sown MPD) on complementarity. Colored lines with shaded 95% confidence intervals represent model predictions from linear mixed-effects models.

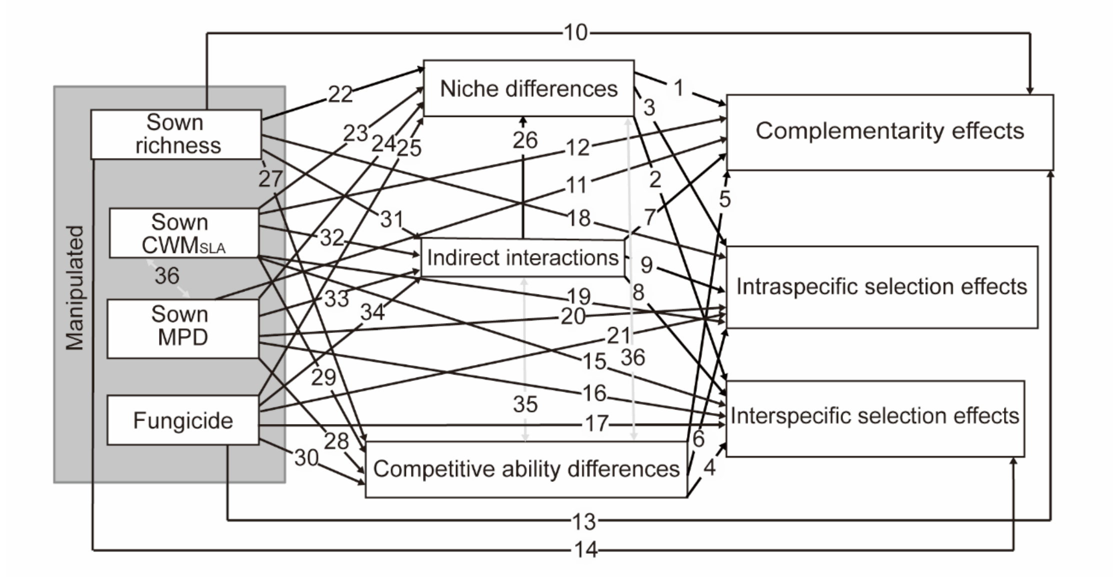


**Fig S4.** The full hypothesized structural equation model linking coexistence mechanisms in multispecies communities to biodiversity effects on productivity across environmental and functional trait contexts. We tested paths from sown species richness (sown richness), functional composition (sown $CWM_{SLA}$), functional diversity (sown MPD), and fungicide application (leaf box) to coexistence mechanisms estimated based on sown species composition to mean biodiversity effects. Coexistence mechanisms comprised standardized niche differences, competitive ability differences, and the contribution of indirect interactions. Biodiversity effects comprised complementarity, interspecific and intraspecific selection effects. Single headed arrows represent hypothesized directional relationships (e.g. path 1 illustrates the effect of niche differences on complementarity effects). Double headed arrows represent covariances, allowing associations between variables without specifying a direction of influence. The numbered paths are explained as follows: 1) Communities with greater niche differences will have higher complementarity effects, due to niche differentiation in resources, enemies and/or abiotic requirement (path 1)[1–3]. 2) Higher niche differences will reduce competitive dominance and weaken interspecific selection (dominance) effects (path 2)[4,5]. 3) Negative intraspecific selection effects (senus Fox's trait-dependent complementarity) arise because some species benefit disproportionally from reduced conspecific limitation in mixtures, communities with greater niche differences will have more negative intraspecific selection effects (path 3)[6]. 4) Higher competitive ability differences will favor dominance by particular species[7,8]. The resulting interspecific selection (dominance) effects may be positive or negative, depending on whether species that dominate have higher or lower monoculture yields (path 4). 5) Higher competitive ability difference will reduce complementarity when dominance suppresses the contributions of subordinate species (path 5)[4]. They may also alter intraspecific selection effects (path 6). 6) Indirect interactions can expand, constrain or have no effect on conditions permitting coexistence, their contribution would be linked to niche differences (path 26)[9,10]. It remains unclear how indirect interactions influence biodiversity effects, we also test direct relationships with complementarity, interspecific and intraspecific selection effects without specifying their directions (path 7, 8 and 9). 7) Higher sown species richness and functional diversity (sown MPD) will lead to higher complementarity (path 10 and 11) and lower interspecific selection effects (path 14 and 16), as many biodiversity-functioning experiments suggested that this could be due to higher niche differentiation (path 22 and 24) and/or lower competitive ability differences (path 27 and 28)[11]. Communities dominated by fast-growing species, indicated by higher sown $CWM_{SLA}$, are expected to exhibit lower complementarity (path 12) and higher interspecific selection effects (path 15)[12]. These relationships could arise through higher competitive ability differences (path 29) and lower niche differences (path 23), for example where inequalities in light competition favor a subset of species[13,14]. 8) Removal of foliar fungal pathogens can erode coexistence by weakening negative density dependence (path 25) and altering competitive hierarchies (path 30)[15,16]. These changes could reduce complementarity (path 13) and/or increase interspecific selection effects when fungal suppression favors species with high monoculture yields (path 17)[17]. 9) Sown species richness, functional diversity, functional composition and fungicide application may influence species' functioning (yield) per unit area in mixtures. We therefore included direct paths from these predictors to intraspecific selection effects without specifying direction (path 18, 19, 20 and 21)[6,18]. 10) Sown species richness, functional diversity, functional composition and fungicide application may influence the contribution of indirections by changing the structure of species interaction networks (path 31, 32, 33 and 34)[19]. These paths were included to test potential drivers for indirect interactions.

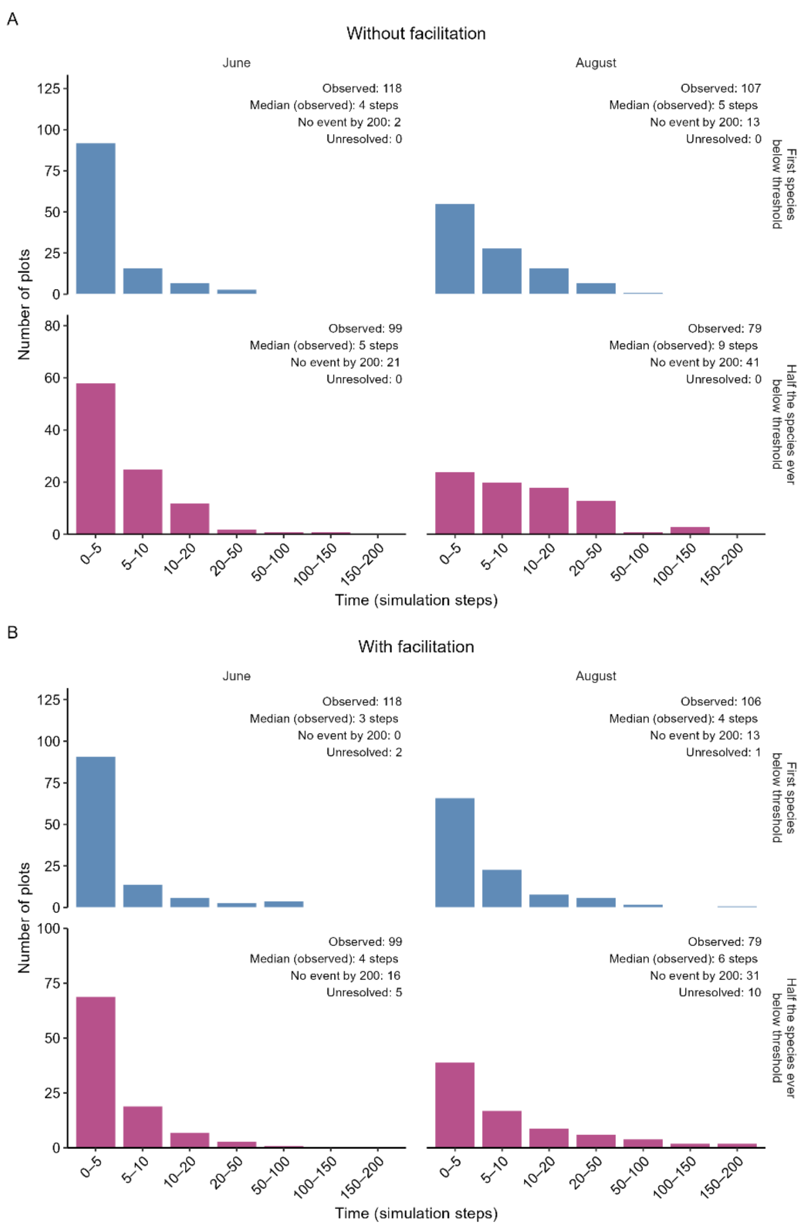


**Fig S5**. Time to species loss in simulated communities with and without facilitation. Distributions of simulation steps until the first species loss and the loss of half the initially sown species are shown separately for June and August under two scenarios: (A) without facilitation and (B) with facilitation. We simulated species abundances for up to 200 steps using a multispecies Ricker model parameterized with interaction coefficients and intrinsic growth rates estimated for each of four treatments and two seasons. Species were considered extinct when their abundance fell below 0.1% of their initial abundances and were subsequently held at zero. Bars show observed events only, meaning that plots without any species loss up to 200 steps were not shown. Annotations report the number of plots with observed events, median event time to those events, the number of simulations that completed 200 steps without the event occurring (the even was not observed until the 200 steps), and the number that stopped before the endpoint could be determined ("unsolved") because a species abundance exceeded the numerical range when facilitation was included. When facilitation was included, 30 and 41 plots stopped before 200 steps because simulated abundances exceeded the numerical range in R. Events recorded before these failures were retained. Relationships between standardized niche differences and time-steps of first species loss or loss of at least half the sown species loss are reported in Table.S4.

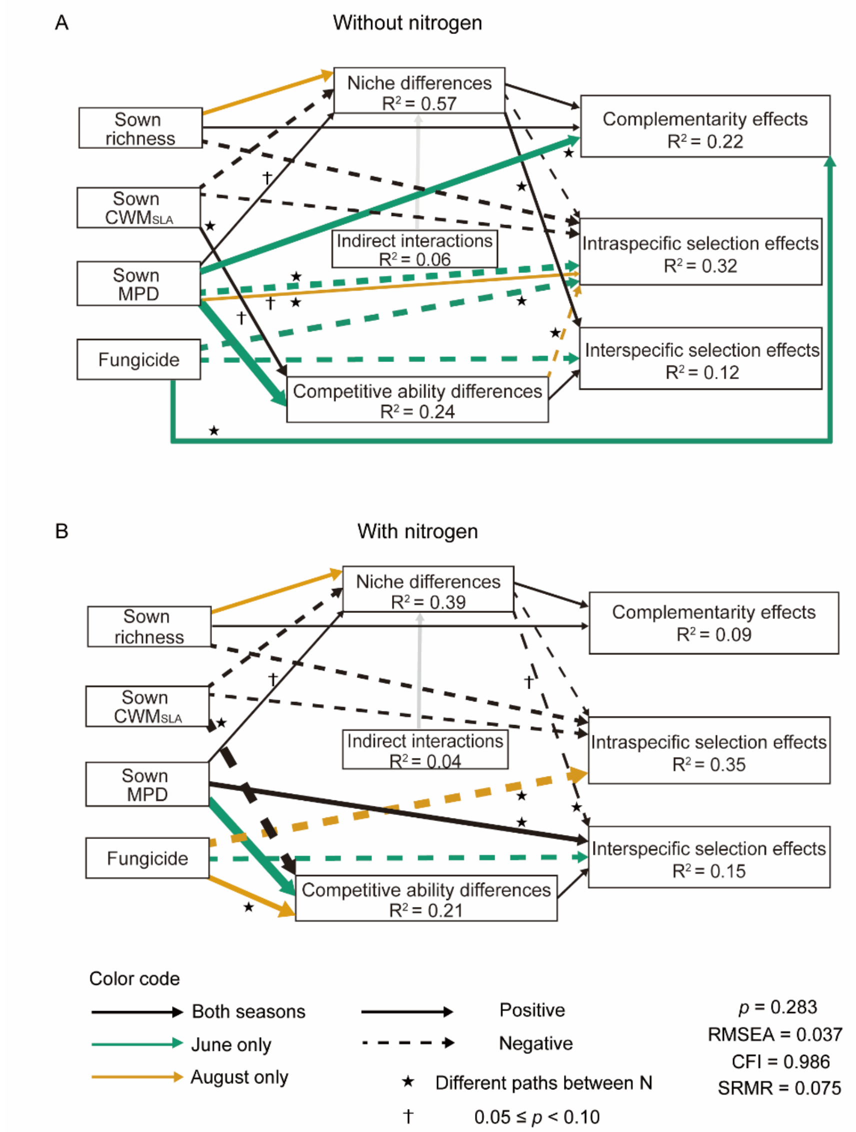


**Fig S6**. Structural equation model linking sown species richness, functional composition (Sown $CWM_{SLA}$), functional diversity (Sown MPD), fungicide, coexistence mechanisms (standardized niche differences, competitive ability differences, and indirect interactions) without facilitation based on sown species composition, and mean diversity effects (Complementarity effects, interspecific and intraspecific selection effects) without nitrogen (A) and with nitrogen (B). Solid and dashed arrows indicate positive and negative effects, respectively. Marginally significant paths $0.05 \leqslant p < 0.10$ are marked with a dagger (†). Paths that are consistent across June and August are shown in black, season-specific paths are highlighted in in bluish-green (June only) or golden-orange (August only). Although niche differences and indirect interactions are mathematically coupled in the structuralist approach, indirect interactions can theoretically promote, constrain or have no effect on coexistence[9]. Based on this theoretical rationale, the directional path from the contribution of indirect interactions to niche differences is shown in grey. Arrow thickness is proportional to the strength of the path. Black stars indicate paths that differ significantly between nitrogen treatments. Non-significant paths ($p \geqslant 0.10$), covariances, and path coefficients are omitted for clarity; full model details are provided in Table S2. $R^2$ values are shown for all endogenous variables, representing the maximum observed across two seasons. Model fit was adequate: $p = 0.283$, CFI = 0.986, RMSEA = 0.037, SRMR = 0.075.

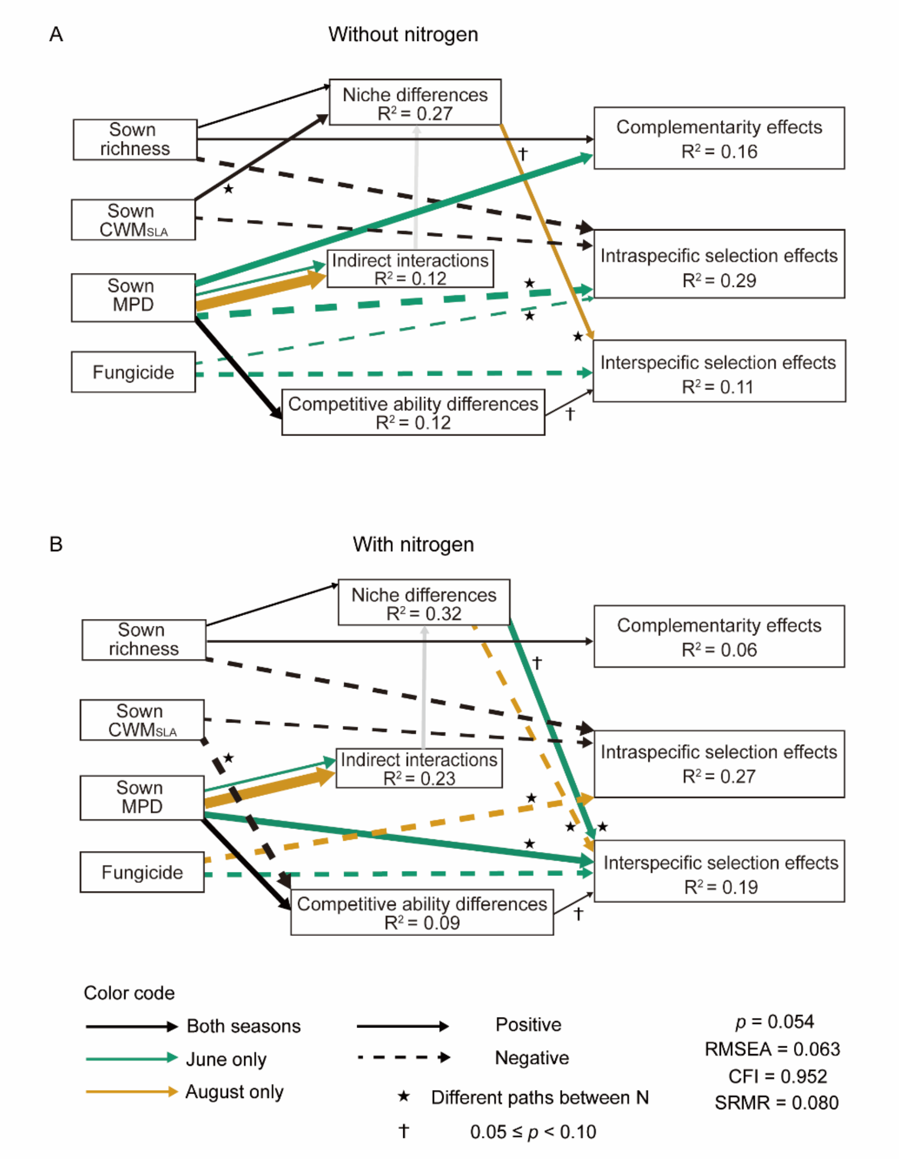


**Fig S7**. Structural equation model linking sown species richness, functional composition (Sown $CWM_{SLA}$), functional diversity (Sown MPD), fungicide, coexistence mechanisms (standardized niche differences, competitive ability differences, and indirect interactions) with facilitation based on sown species composition, and mean diversity effects (Complementarity effects, interspecific and intraspecific selection effects) without nitrogen (A) and with nitrogen (B). Solid and dashed arrows indicate positive and negative effects, respectively. Marginally significant paths $0.05 \leqslant p < 0.10$ are marked with a dagger (†). Paths that are consistent across June and August are shown in black, season-specific paths are highlighted in in bluish-green (June only) or golden-orange (August only). Although niche differences and indirect interactions are mathematically coupled in the structuralist approach, indirect interactions can theoretically promote, constrain or have no effect on coexistence[9]. Based on this theoretical rationale, we the directional path from the contribution of indirect interactions to niche differences is shown in grey. Arrow thickness is proportional to the strength of the path. Black stars indicate paths that differ significantly between nitrogen treatments. Non-significant paths ($p \geqslant 0.10$), covariances, and path coefficients are omitted for clarity; full model details are provided in Table S2. $R^2$ values are shown for all endogenous variables, representing the maximum observed across two seasons. Model fit was adequate: $p = 0.054$, CFI = 0.952, RMSEA = 0.063, SRMR = 0.080.

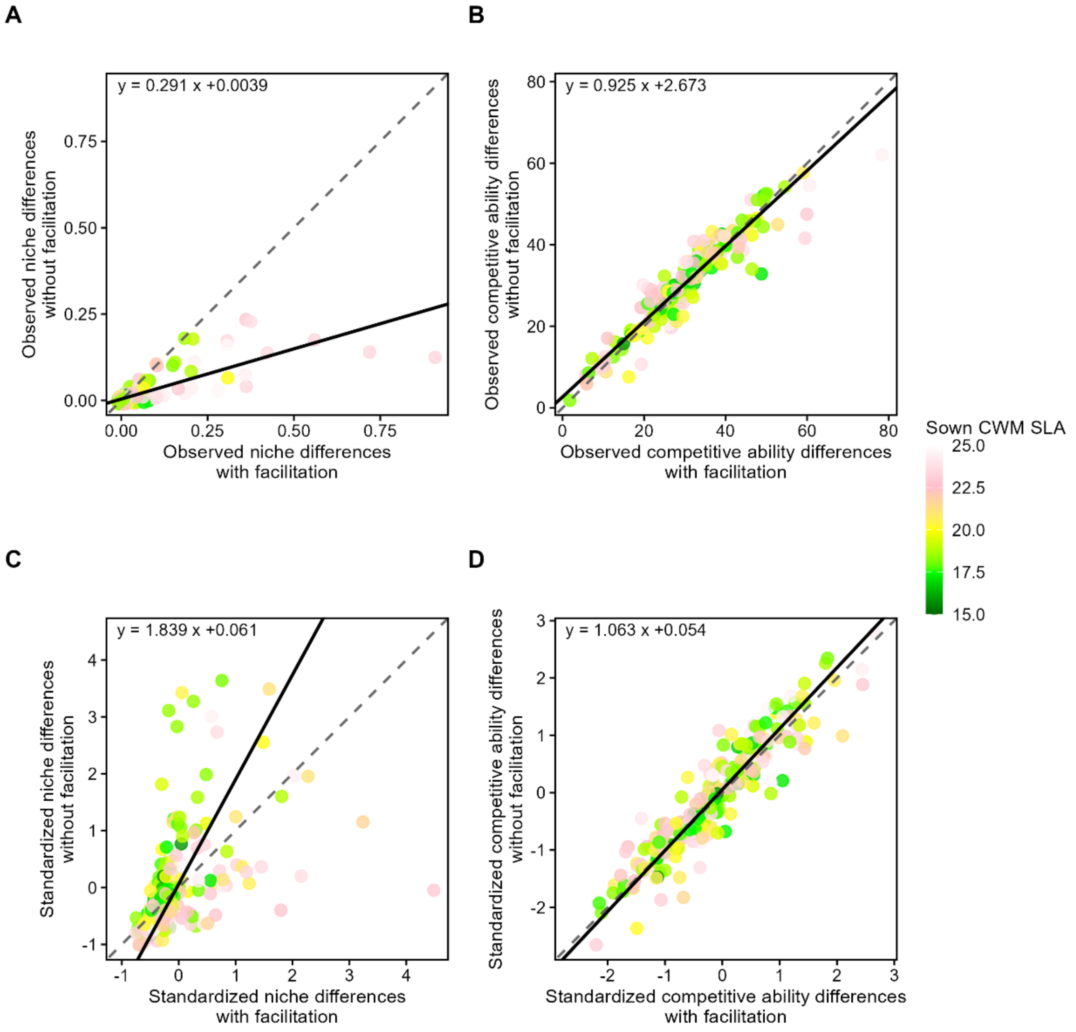


**Fig S8**. Relationships between structural niche differences and competitive ability differences calculated with and without facilitation. Point colors indicate sown community-weighted mean of specific leaf area (sown $CWM_{SLA}$). (A) Observed niche differences with and without facilitation ($R^2 = 0.60$, $p < 0.001$). (B) Observed competitive ability differences with and without facilitation ($R^2 = 0.88$, $p < 0.001$). (C) Standardized niche differences with and without facilitation ($R^2 = 0.23$, $p < 0.001$). (D) Standardized competitive ability differences with and without facilitation ($R^2 = 0.86$, $p < 0.001$). Dashed lines represent the 1:1 relationship, and the solid lines show fitted orthogonal regressions. The fitted equation is displayed in each panel, where x and y represent values calculated with and without facilitation, respectively. In (A) and (B), observed niche and competitive ability differences were estimated based on the sown species composition. In (C) and (D), we applied a randomization by shuffling species identifies in the matrices of intrinsic growth rates and competition coefficients, under a given treatment and season. We ensured species identities remained consistent between the two matrices, which preserved the inherent connection between competition coefficients and intrinsic growth rates. We used the randomized matrices (99 times) and recalculated the expected niche and competitive ability differences. The standardized niche and competitive ability differences were calculated using the observed values of the niche and competitive ability differences standardized by the mean expected niche and competitive ability differences for each community.

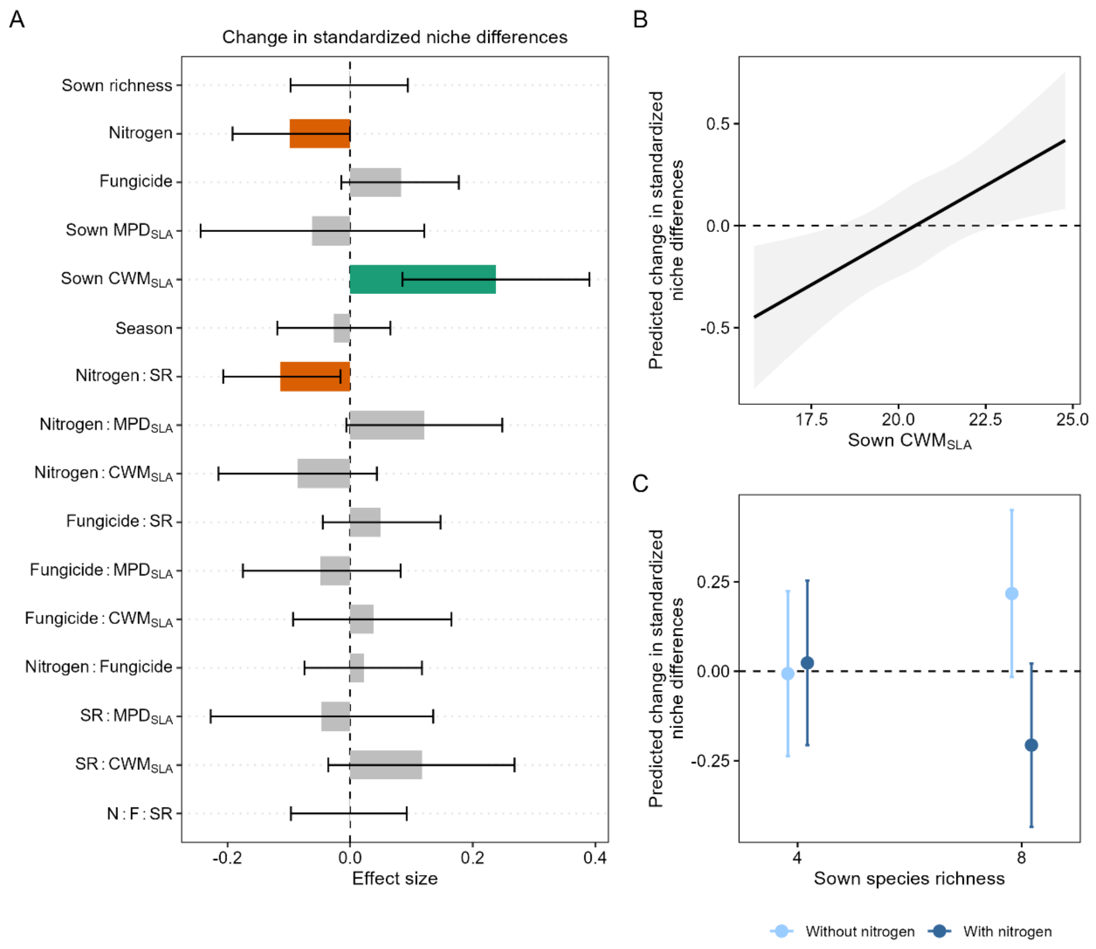


**Fig S9**. Results of linear mixed effect models showing drivers of the difference in standardized niche differences with and without facilitation. (A), Shown are mean standardized effects (± confidence intervals) of each scaled and centered variable on the difference in standardized niche differences with and without facilitation. Positive effects are shown in green, negative effects in orange, and non-significant effects are in gray. SR: sown species richness, Sown MPD: mean pairwise distance in SLA based on sown composition, Sown $CWM_{SLA}$: community weighted mean of SLA based on sown composition, N: nitrogen, F: fungicide. (B) Direct effects of functional composition ($CWM_{SLA}$). (C) Interaction between sown species richness and nitrogen addition.

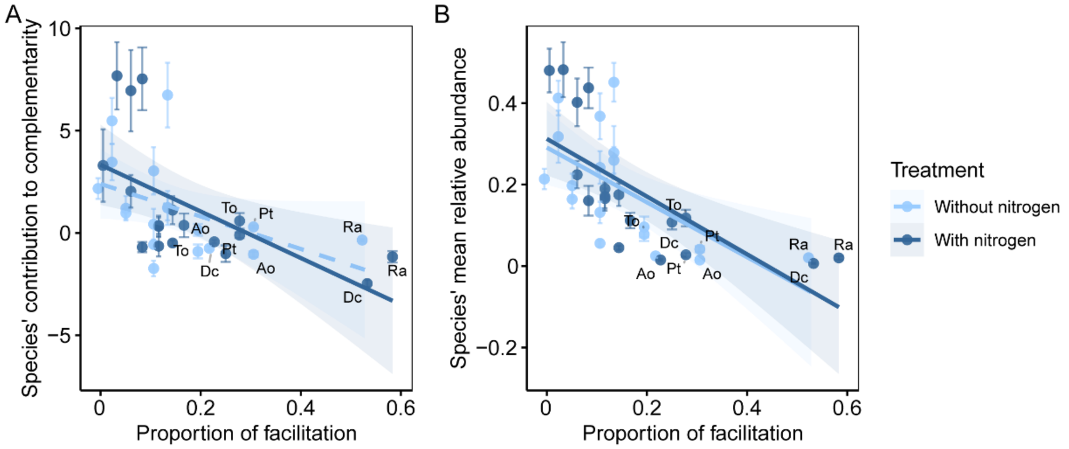


**Fig S10.** Relationships between species' contributions to complementarity effects ($\Delta RY_i \times M_i$) and the proportion of facilitation each species received (A), and between species' mean relative abundance and the proportion of facilitation received (B) in August. Solid lines represent marginally significant ($p < 0.\ 05$) fit from linear regressions across treatments with shaded areas represent 95% confidence intervals. To improve readability, only the top five species receiving the highest levels of facilitation are labeled. Species abbreviations are listed here: Ra: *Rumex acetosa*; Ao: *Athoxanthum odoratum*; Dc: *Daucus carota*; To: *Taraxacum officinale*; Pt: *Poa trivialis*. In (A), species that received more facilitation had lower complementarity effects (With nitrogen: $p < 0.01$, $R^2 = 0.31$; Without nitrogen: $p = 0.06$, $R^2 = 0.15$). In (B), species that received more facilitation had lower relative abundances (With nitrogen: $p < 0.01$, $R^2 = 0.47$; Without nitrogen: $p < 0.01$, $R^2 = 0.34$).

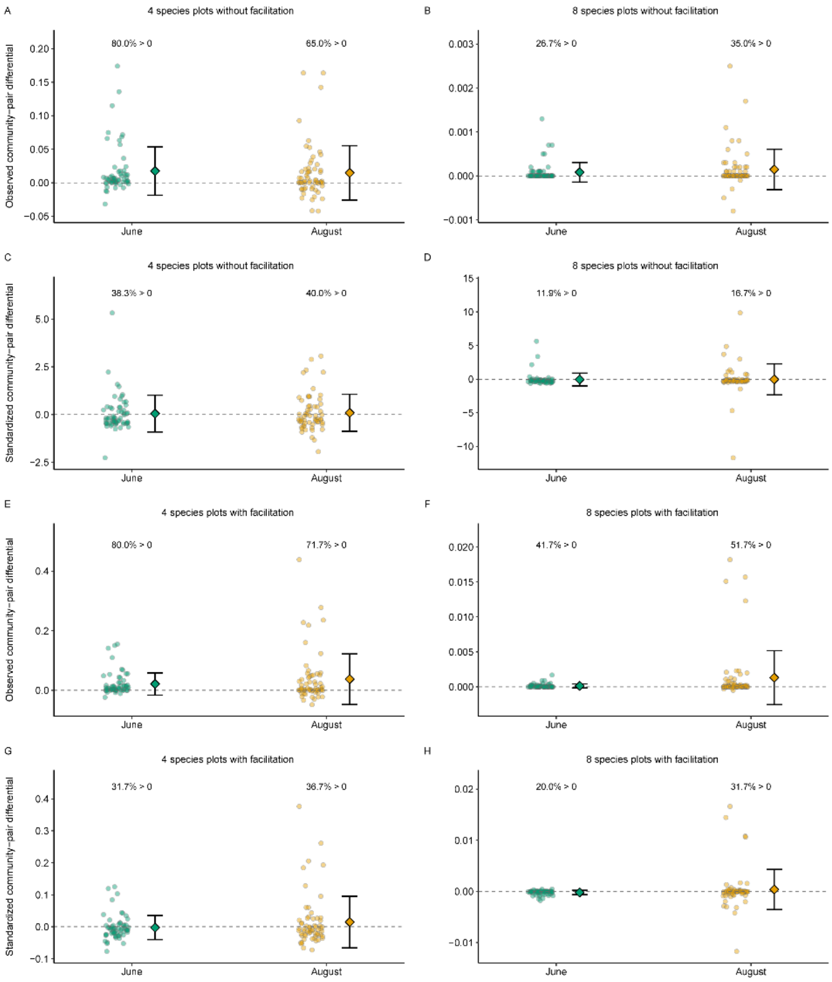


**Fig S11.** Observed and standardized community-pair differential (the contribution of indirect interactions to coexistence) in four- and eight-species communities, with and without facilitation. A-D show results without facilitation, and E-H show results with facilitation. Observed community-pair differentials are shown for four-species (A, E) and eight-species communities (B, F); standardized community-pair differentials are shown for four-species (C, G) and eight-species communities (D, H). The observed community-pair differential, D, is the difference between the full-community feasibility domain and the intersection of pairwise feasibility domains. Positive and negative observed values indicate that indirect interactions expand or reduce opportunities for coexistence, respectively. Standardized values express the deviation of observed D from its randomization mean in units of the randomization standard deviation; their signs therefore indicate deviations from the null expectation. Circles represent individual communities, and diamonds and error bars indicate means ±SD. Colors distinguish between June (green) and August (orange). Percentages indicate the proportion of communities with values above zero within each season and panel.

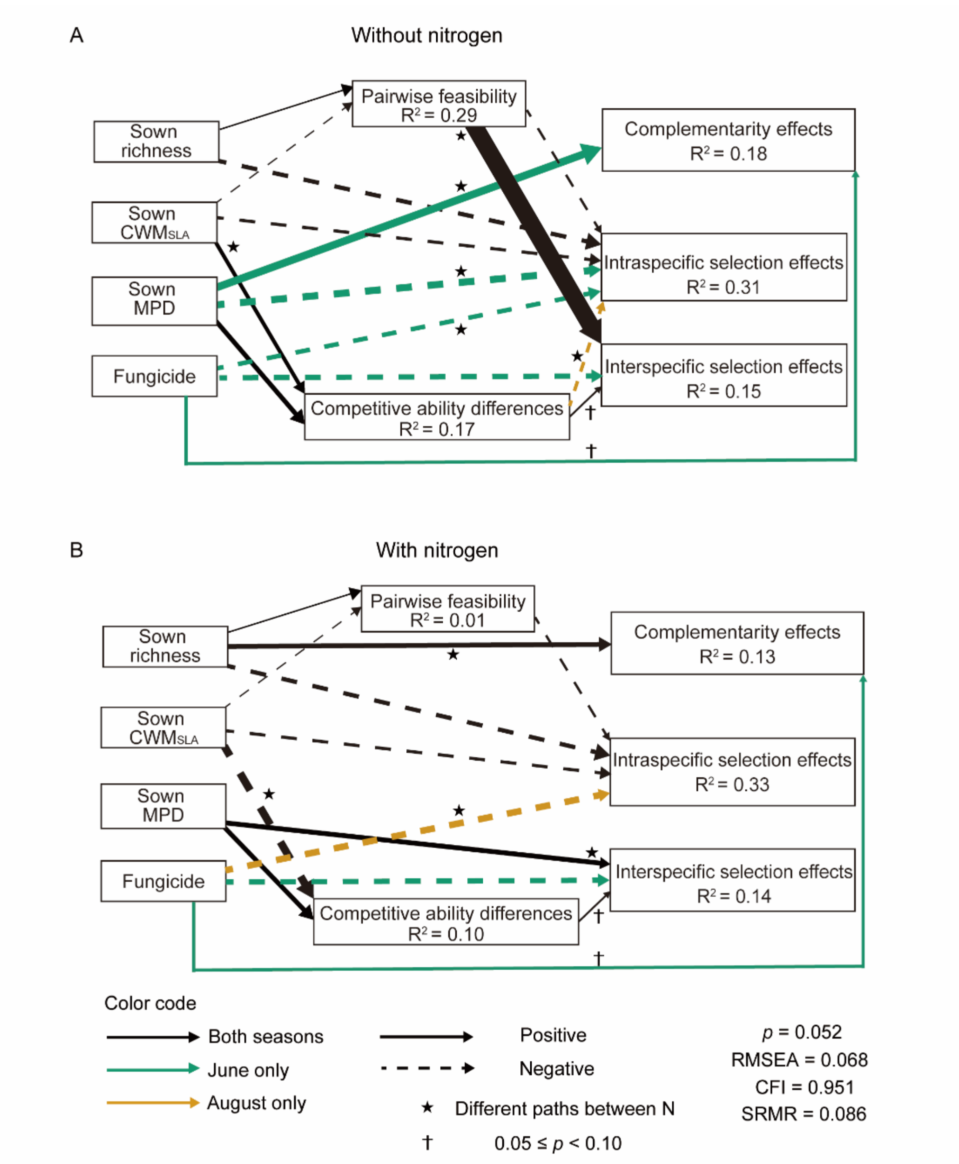


**Fig S12**. Structural equation model linking sown species richness, functional composition (Sown $CWM_{SLA}$), functional diversity (Sown MPD), fungicide, standardized niche differences for all species pairs and standardized competitive ability differences without facilitation based on sown species composition, and mean diversity effects (Complementarity effects, interspecific and intraspecific selection effects) without nitrogen (A) and with nitrogen (B). To test whether standardized niche differences for all species pairs (the feasibility domain for all pairs) are sufficient to promote complementarity effects, we directly used the feasibility domain for all pairs without considering the contribution of indirect interactions in the model. Solid and dashed arrows indicate positive and negative effects, respectively. Solid and dashed arrows indicate positive and negative effects, respectively. Marginally significant paths $0.05 \leqslant p < 0.10$ are marked with a dagger (†). Paths that are consistent across June and August are shown in black, season-specific paths are highlighted in in bluish-green (June only) or golden-orange (August only). Arrow thickness is proportional to the strength of the path. Black stars indicate paths that differ significantly between nitrogen treatments. Non-significant paths ($p \geqslant 0.10$), covariances, and path coefficients are omitted for clarity; full model details are provided in Table S4. $R^2$ values are shown for all endogenous variables, representing the maximum observed across two seasons. Model fit measures were: $p = 0.052$, CFI = 0.951, RMSEA = 0.068, SRMR = 0.086.

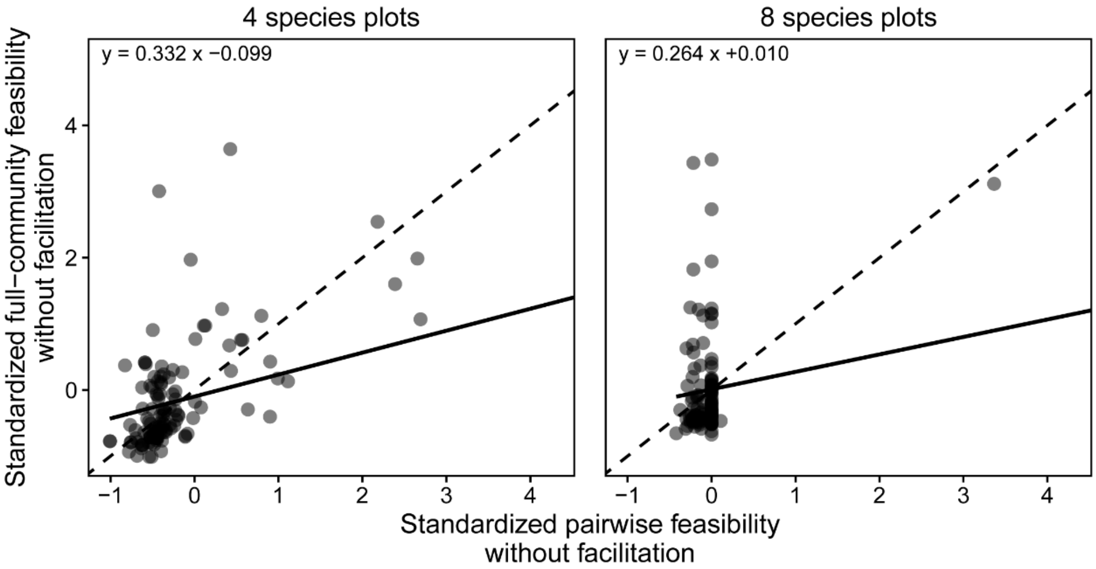


**Fig S13**. Relationships between standardized pairwise feasibility (Ω_all) and standardized full-community feasibility (standardized niche differences, Ω) in 4- and 8-species plots without facilitation. Each point represents one plot per season. Dashed lines represent the 1:1 relationship, and the solid lines show fitted orthogonal regressions. The fitted equation is displayed in each panel for each richness level, where x and y represent values of standardized pairwise feasibility and standardized full-community feasibility without facilitation, respectively. In 8-species plots, the pairwise feasibility (Ω_all) values clustered near zero, suggesting indirection interactions are important in more diverse plots for standardized full-community feasibility (standardized niche differences, Ω). In both panels, to improve the visibility, the x-axis is displayed up to the $99^{th}$ percentile of standardized pairwise feasibility.

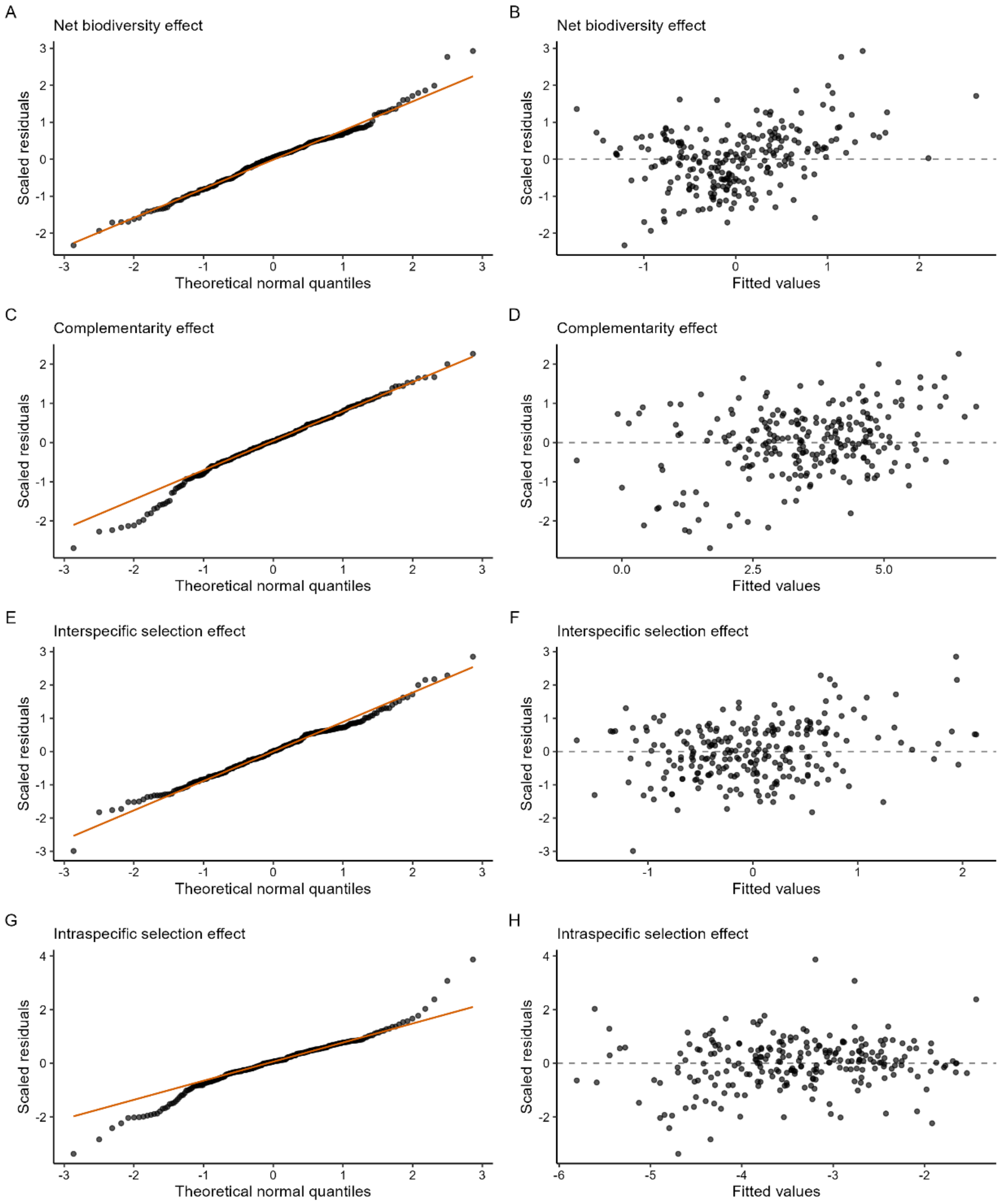


**Fig S14.** Residual diagnostics for the linear mixed-effects models of biodiversity effects. Normal quantile-quantile plots (Q-Q plots, left column) and residual-versus-fitted plots (right column) are shown for net biodiversity effects (A-B), complementarity effects (C-D), interspecific selection effects (E-F), and intraspecific selection effects (G-H). Orange lines indicate normal-reference lines in the Q-Q plots; dashed horizontal lines indicate zero residuals. Fitted values are shown on the response scale used in each model.

**Table S1**. Results of linear mixed effect models testing the effects of sown species richness, functional composition (Sown $CWM_{SLA}$) and functional diversity (Sown MPD), treatments (Nitrogen × Fungicide) and their interactions on mean biodiversity effects per plot across two seasons.

| | Net diversity effect | | | Complementarity effects | | | Intraspecific selection effects | | | Interspecific selection effects | | |
|---|---|---|---|---|---|---|---|---|---|---|---|---|
| | df | F-value | *p* | df | F-value | *p* | df | F-value | *P* | df | F-value | *p* |
| N | 1, 80.30 | 0.86 | 0.36 | 1, 80.23 | 0.73 | 0.40 | 1, 80.60 | 0.79 | 0.38 | 1, 81.11 | 29.58 | **< 0.001** |
| F | 1, 84.22 | 1.43 | 0.23 | 1, 83.29 | 1.29 | 0.26 | 1, 82.62 | 3.45 | **0.07** | 1, 87.53 | 2.42 | 0.12 |
| SR | 1, 24.13 | 0.05 | 0.83 | 1, 24.09 | 3.84 | **0.06** | 1, 24.05 | 13.61 | **0.001** | 1, 24.26 | 0.02 | 0.88 |
| Sown $CWM_{SLA}$ | 1, 30.56 | 0.40 | 0.53 | 1, 29.10 | 0.001 | 0.97 | 1, 27.35 | 5.03 | **0.03** | 1, 36.43 | 0.02 | 0.89 |
| Sown MPD | 1, 32.38 | 1.01 | 0.32 | 1, 30.32 | 0.86 | 0.36 | 1, 27.85 | 0.42 | 0.52 | 1, 41.65 | 0.27 | 0.61 |
| Season | 1, 119 | 35.79 | **< 0.001** | 1, 119 | 32.31 | **< 0.001** | 1, 119 | 39.47 | **< 0.001** | 1, 119 | 3.86 | **0.05** |
| N × F | 1, 82.07 | 0.43 | 0.51 | 1, 81.31 | 0.01 | 0.92 | 1, 81.91 | 0.27 | 0.760 | 1, 85.30 | 0.83 | 0.37 |
| N × SR | 1, 79.95 | 3.35 | **0.07** | 1, 79.85 | 6.43 | **0.01** | 1, 80.45 | 3.82 | **0.05** | 1, 81.12 | 0.004 | 0.95 |
| N × Sown $CWM_{SLA}$ | 1, 82.58 | 0.05 | 0.82 | 1, 82.33 | 0.74 | 0.39 | 1, 82.07 | 3.13 | **0.08** | 1, 83.30 | 0.44 | 0.51 |
| N × Sown MPD | 1, 80.27 | 0.95 | 0.33 | 1, 80.55 | 5.87 | **0.02** | 1, 81.12 | 2.97 | **0.09** | 1, 79.85 | 1.54 | 0.22 |
| F × SR | 1, 86.21 | 0.43 | 0.51 | 1, 85.24 | 0.38 | 0.54 | 1, 80.42 | 0.14 | 0.71 | 1, 89.73 | 0.0002 | 0.99 |
| F × Sown $CWM_{SLA}$ | 1, 80.07 | 2.11 | 0.15 | 1, 80.61 | 0.07 | 0.80 | 1, 82.48 | 0.35 | 0.56 | 1, 79.94 | 1.57 | 0.21 |
| F × Sown MPD | 1, 81.68 | 1.64 | 0.20 | 1, 82.17 | 0.12 | 0.73 | 1, 84.13 | 0.08 | 0.77 | 1, 81.76 | 1.28 | 0.26 |
| SR × Sown $CWM_{SLA}$ | 1, 31.24 | 0.41 | 0.53 | 1, 29.54 | 0.18 | 0.67 | 1, 27.44 | 0.20 | 0.65 | 1, 37.64 | 0.01 | 0.93 |
| SR × Sown MPD | 1, 33.66 | 1.37 | 0.25 | 1, 31.32 | 0.14 | 0.71 | 1, 28.54 | 0.56 | 0.46 | 1, 43.14 | 0.19 | 0.67 |
| N × F × SR | 1, 83.29 | 2.61 | 0.11 | 1, 82.49 | 4.04 | **0.05** | 1, 82.15 | 0.77 | 0.38 | 1, 86.72 | 2.77 | 0.10 |

**Table S4**. Results of linear mixed effect models testing the effects of standardized niche differences and seasons on time-steps of first species loss or loss of at least half the sown species per plot.

| Facilitation | Time-steps of the event | Fixed effect | Estimate ± SE | df | F-value | *p* |
|---|---|---|---|---|---|---|
| Without | The first species loss | Standardized niche differences | **0.16 ± 0.06** | **1, 229.36** | **6.23** | **0.01** |
| | | Season | **0.35 ± 0.05** | **1, 118.20** | **45.14** | **< 0.001** |
| | Loss of at least half the sown species | Standardized niche differences | **0.21 ± 0.09** | **1, 232.66** | **4.64** | **0.03** |
| | | Season | **0.46 ± 0.08** | **1, 118.22** | **32.85** | **< 0.001** |
| With | The first species loss | Standardized niche differences | 0.04 ± 0.07 | 1, 230.65 | 0.37 | 0.54 |
| | | Season | **0.32 ± 0.06** | **1, 117.29** | **29.17** | **< 0.001** |
| | Loss of at least half the sown species | Standardized niche differences | -0.06 ± 0.10 | 1, 220.22 | 0.40 | 0.53 |
| | | Season | **0.43 ± 0.09** | **1, 113.02** | **23.50** | **< 0.001** |

**Table S3**. Structural equation model (SEM) results. Coexistence mechanisms were quantified with and without facilitation. Although niche differences and indirect interactions are mathematically coupled in the structuralist approach, indirect interactions can theoretically promote, constrain or have no effect on coexistence[9]. Based on this theoretical rationale, we specified a directional path from the contribution of indirect interactions to niche differences, rather than modelling their association as a covariance. Each row reports one path. To reduce the redundancy, if a path was constrained across four groups (N × season), only the parameter estimates without nitrogen in June are shown (as it is the same in other groups). If a path differed between nitrogen treatments but was constrained between two seasons, only the parameter estimates in June are shown separately for with and without nitrogen. If a path differed between seasons but was constrained between nitrogen treatments within each season, only without nitrogen estimates are shown separately for June and August. And if a path could not be constrained (i.e., it differed among groups), estimates for all four groups are shown. Model fit was adequate: SEM without facilitation: $p = 0.283$, CFI = 0.986, RMSEA = 0.037, SRMR = 0.075; SEM with facilitation: $p = 0.054$, CFI = 0.952, RMSEA = 0.063, SRMR = 0.080.

| Response | Predictor | Treatment | Constrain | Coefficient | SE | *P* value |
|---|---|---|---|---|---|---|
| **Coexistence mechanisms estimated without facilitation** | | | | | | |
| Complementarity effects | Niche differences | June & without N | All | **0.14** | **0.05** | **0.01** |
| | Competitive ability differences | June & without N | All | -0.04 | 0.06 | 0.45 |
| | Indirect effects | June & without N | All | 0.001 | 0.06 | 0.98 |
| | Sown richness | June & without N | All | **0.12** | **0.05** | **0.01** |
| | Sown MPD | June & without N | None | **0.35** | **0.13** | **0.006** |
| | | June & with N | | -0.20 | 0.15 | 0.17 |
| | | August & without N | | 0.09 | 0.09 | 0.42 |
| | | August & with N | | 0.08 | 0.11 | 0.51 |
| | Sown $CWM_{SLA}$ | June & without N | All | 0.01 | 0.07 | 0.91 |
| | Fungicide | June & without N | None | **0.28** | **0.12** | **0.02** |
| | | June & with N | | 0.04 | 0.15 | 0.79 |
| | | August & without N | | -0.08 | 0.08 | 0.27 |
| | | August & with N | | 0.13 | 0.11 | 0.23 |
| Interspecific | Niche | With N | N | **-0.15** | **0.08** | **0.06** |

| | | | | | | |
|---|---|---|---|---|---|---|
| selection effects | differences | Without N | | **0.16** | **0.07** | **0.03** |
| | Competitive ability differences | June & without N | All | **0.12** | **0.05** | **0.02** |
| | Indirect effects | June & without N | All | -0.04 | 0.06 | 0.46 |
| | Sown richness | Competitive ability differences | June & without N | 0.03 | 0.05 | 0.56 |
| | Sown MPD | With N | N | **-0.27** | **0.09** | **0.004** |
| | | Without N | | -0.05 | 0.08 | 0.52 |
| | Sown $CWM_{SLA}$ | June & without N | All | 0.02 | 0.06 | 0.71 |
| | Fungicide | June | Season | **-0.27** | **0.09** | **0.003** |
| | | August | | 0.05 | 0.06 | 0.41 |
| Intraspecific selection effects | Niche differences | June & without N | All | **-0.12** | **0.05** | **0.02** |
| | Competitive ability differences | June & without N | None | 0.09 | 0.07 | 0.20 |
| | | June & with N | | 0.17 | 0.11 | 0.13 |
| | | August & without N | | **-0.17** | **0.08** | **0.04** |
| | | August & with N | | 0.10 | 0.10 | 0.32 |
| | Indirect effects | June & without N | All | 0.08 | 0.05 | 0.13 |
| | Sown richness | June & without N | All | **-0.22** | **0.05** | **< 0.001** |
| | Sown MPD | June & without N | None | **-0.34** | **0.12** | **0.005** |
| | | June & with N | | 0.01 | 0.12 | 0.93 |
| | | August & without N | | **0.17** | **0.09** | **0.077** |
| | | August & with N | | -0.16 | 0.11 | 0.16 |
| | Sown $CWM_{SLA}$ | June without N | All | **-0.18** | **0.06** | **0.005** |
| | Fungicide | June & without N | None | **-0.32** | **0.11** | **0.003** |

| | | | | | | |
|---|---|---|---|---|---|---|
| | | June & with N | | 0.07 | 0.12 | 0.52 |
| | | August & without N | | 0.10 | 0.08 | 0.22 |
| | | August & with N | | **-0.42** | **0.11** | **< 0.001** |
| Niche differences | Sown richness | June | Season | -0.05 | 0.07 | 0.45 |
| | | August | | **0.21** | **0.06** | **0.001** |
| | Sown MPD | June without N | All | **0.12** | **0.06** | **0.054** |
| | Sown $CWM_{SLA}$ | June & without N | All | **-0.21** | **0.06** | **0.001** |
| | Indirect effects | June & without N | None | **0.77** | **0.17** | **< 0.001** |
| | | June & with N | | **0.84** | **0.14** | **< 0.001** |
| | | August & without N | | **1.08** | **0.13** | **< 0.001** |
| | | August & with N | | **-0.20** | **0.08** | **0. 02** |
| | Fungicide | June & without N | All | -0.02 | 0.05 | 0.74 |
| Competitive ability differences | Sown richness | June & without N | All | 0.02 | 0.06 | 0.69 |
| | Sown MPD | June | Season | **0.48** | **0.09** | **< 0.001** |
| | | August | | 0.12 | 0.09 | 0.19 |
| | Sown $CWM_{SLA}$ | With N | N | **-0.46** | **0.09** | **< 0.001** |
| | | Without N | | **0.17** | **0.10** | **0.08** |
| | Fungicide | June & without N | None | -0.05 | 0.13 | 0.71 |
| | | June & with N | | -0.08 | 0.10 | 0.41 |
| | | August & without N | | -0.12 | 0.11 | 0.27 |
| | | August & with N | | **0.28** | **0.11** | **0.02** |
| The contribution of indirect interactions | Sown richness | June & without N | All | -0.05 | 0.05 | 0.26 |
| | Sown MPD | June & without N | All | 0.08 | 0.06 | 0.17 |
| | Sown $CWM_{SLA}$ | June & without N | All | 0.06 | 0.06 | 0.32 |
| | Fungicide | June & | All | -0.03 | 0.05 | 0.51 |

| | | | | | | |
|---|---|---|---|---|---|---|
| | | without N | | | | |
| **Coexistence mechanisms estimated with facilitation** | | | | | | |
| Complementarity effects | Niche differences | June & without N | All | 0.07 | 0.06 | 0.22 |
| | Competitive ability differences | June & without N | All | 0.02 | 0.05 | 0.70 |
| | Indirect interactions | June & without N | All | -0.01 | 0.05 | 0.86 |
| | Sown richness | June & without N | All | **0.13** | **0.05** | **0.01** |
| | Sown MPD | June & without N | None | **0.36** | **0.13** | **0.007** |
| | | June & with N | | -0.21 | 0.15 | 0.15 |
| | | August & without N | | 0.07 | 0.10 | 0.48 |
| | | August & with N | | 0.07 | 0.12 | 0.55 |
| | Sown $CWM_{SLA}$ | June & without N | All | -0.01 | 0.07 | 0.87 |
| | Fungicide | June & without N | All | 0.05 | 0.05 | 0.36 |
| Interspecific selection effects | Niche differences | June & without N | None | 0.11 | 0.08 | 0.16 |
| | | June & with N | | **0.33** | **0.20** | **0.10** |
| | | August & without N | | **0.21** | **0.12** | **0.08** |
| | | August & with N | | **-0.26** | **0.12** | **0.03** |
| | Competitive ability differences | June & without N | All | **0.09** | **0.05** | **0.07** |
| | Indirect interactions | June & without N | All | -0.05 | 0.05 | 0.25 |
| | Sown richness | June & without N | All | 0.01 | 0.05 | 0.78 |
| | Sown MPD | With N | N | **0.35** | **0.09** | **< 0.001** |
| | | Without N | | -0.04 | 0.08 | 0.66 |
| | Sown $CWM_{SLA}$ | June & without N | All | 0.003 | 0.06 | 0.96 |
| | Fungicide | June | Season | **-0.26** | **0.09** | **0.004** |

| | | | | | | |
|---|---|---|---|---|---|---|
| | | August | | 0.06 | 0.06 | 0.26 |
| Intraspecific selection effects | Niche differences | June & without N | All | 0.06 | 0.06 | 0.27 |
| | Competitive ability differences | June & without N | All | -0.03 | 0.05 | 0.63 |
| | Indirect interactions | June & without N | All | -0.02 | 0.05 | 0.69 |
| | Sown richness | June & without N | All | **-0.24** | **0.05** | **< 0.001** |
| | Sown MPD | June & without N | None | **-0.36** | **0.12** | **0.003** |
| | | June & with N | | 0.07 | 0.12 | 0.58 |
| | | August & without N | | 0.12 | 0.10 | 0.21 |
| | | August & with N | | -0.16 | 0.12 | 0.18 |
| | Sown $CWM_{SLA}$ | June & without N | All | **-0.18** | **0.06** | **0.005** |
| | Fungicide | June & without N | None | **-0.15** | **0.07** | **0.04** |
| | | June & with N | | 0.06 | 0.10 | 0.54 |
| | | August & without N | | 0.04 | 0.07 | 0.63 |
| | | August & with N | | **-0.31** | **0.10** | **0.001** |
| Niche differences | Sown richness | June & without N | All | **0.11** | **0.05** | **0.02** |
| | Sown MPD | June & without N | All | 0.08 | 0.07 | 0.23 |
| | Sown $CWM_{SLA}$ | With N | N | -0.08 | 0.08 | 0.32 |
| | | Without N | | **0.19** | **0.08** | **0.01** |
| | Indirect interactions | June & without N | None | **0.70** | **0.25** | **0.005** |
| | | June & with N | | 0.13 | 0.21 | 0.55 |
| | | August & without N | | **0.12** | **0.05** | **0.02** |
| | | August & with N | | **0.44** | **0.09** | **< 0.001** |
| | Fungicide | June & | All | 0.01 | 0.05 | 0.76 |

| | | | | | | |
|---|---|---|---|---|---|---|
| | | without N | | | | |
| Competitive ability differences | Sown richness | June & without N | All | 0 | 0.06 | 0.99 |
| | Sown MPD | June & without N | All | **0.29** | **0.08** | **< 0.001** |
| | Sown $CWM_{SLA}$ | With N | N | **-0.36** | **0.10** | **< 0.001** |
| | | Without N | | 0.10 | 0.10 | 0.33 |
| | Fungicide | June & without N | All | -0.01 | 0.06 | 0.88 |
| Indirect interactions | Sown richness | June & without N | All | -0.004 | 0.04 | 0.92 |
| | Sown MPD | June | Season | **0.14** | **0.06** | **0.02** |
| | | August | | **0.55** | **0.11** | **< 0.001** |
| | Sown $CWM_{SLA}$ | June & without N | All | -0.07 | 0.06 | 0.26 |
| | Fungicide | June & without N | All | 0.03 | 0.04 | 0.54 |

**Table S4**. Structural equation model (SEM) results using the standardized pairwise feasibility domain (Ω_all) in place of community-level standardized niche differences (Ω). Following Saavedra et al.[9], Ω_all quantifies the fraction of intrinsic growth rate space compatible with coexistence of every species pair considered in isolation. We replaced standardized Ω with standardized Ω_all to assess whether relationships with complementarity depend on using full-community feasibility or the conditions permitting coexistence of all species pairs. This was only tested without facilitation, because the positive effect of standardized niche differences on complementarity effects only appeared with facilitation. Each row reports one path. To reduce the redundancy, if a path was constrained across four groups (N × season), only the parameter estimates without nitrogen in June are shown (as it is the same in other groups). If a path differed between nitrogen treatments but was constrained between two seasons, only the parameter estimates in June are shown separately for with and without nitrogen. If a path differed between seasons but was constrained between nitrogen treatments within each season, only without nitrogen estimates are shown separately for June and August. And if a path could not be constrained (i.e., it differed among groups), estimates for all four groups are shown. Model fit was adequate: $p$ = 0.052, CFI =0.951, RMSEA = 0.068, SRMR = 0.086.

| **Response** | **Predictor** | **Treatment** | **Constrain** | **Coefficient** | **SE** | ***P* value** |
|---|---|---|---|---|---|---|
| Complementarity effects | Ω_all | June & without N | All | 0.09 | 0.06 | 0.12 |
| | Competitive ability differences | June & without N | All | -0.03 | 0.05 | 0.59 |
| | Sown richness | With N | N | **0.26** | **0.08** | **0.001** |
| | | Without N | | 0.08 | 0.06 | 0.17 |
| | Sown MPD | June & without N | None | **0.40** | **0.13** | **0.002** |
| | | June & with N | | -0.21 | 0.15 | 0.17 |
| | | August & without N | | 0.10 | 0.09 | 0.29 |
| | | August & with N | | 0.09 | 0.11 | 0.41 |
| | Sown $CWM_{SLA}$ | June & without N | All | -0.005 | 0.07 | 0.93 |
| | Fungicide | June | Season | **0.17** | **0.09** | **0.06** |
| | | August | | -0.01 | 0.06 | 0.92 |
| Interspecific selection effects | Ω_all | With N | N | -0.04 | 0.06 | 0.49 |
| | | Without N | | **0.91** | **0.34** | **0.007** |
| | Competitive ability differences | June & without N | All | **0.10** | **0.05** | **0.06** |
| | Sown | June & | All | -0.02 | 0.05 | 0.64 |

| | | | | | | |
|---|---|---|---|---|---|---|
| | richness | without N | | | | |
| | Sown MPD | With N | N | **0.25** | **0.09** | **0.008** |
| | | Without N | | -0.02 | 0.08 | 0.82 |
| | Sown $CWM_{SLA}$ | June & without N | All | 0.05 | 0.07 | 0.42 |
| | Fungicide | June | Season | **-0.28** | **0.09** | **0.003** |
| | | August | | 0.09 | 0.06 | 0.13 |
| Intraspecific selection effects | Ω_all | June & without N | All | **-0.13** | **0.05** | **0.02** |
| | Competitive ability differences | June & without N | None | 0.08 | 0.07 | 0.30 |
| | | June & with N | | 0.16 | 0.10 | 0.13 |
| | | August & without N | | **-0.18** | **0.08** | **0.03** |
| | | August & with N | | 0.09 | 0.09 | 0.35 |
| | Sown richness | June & without N | All | **-0.23** | **0.05** | **< 0.001** |
| | Sown MPD | June & without N | None | **-0.37** | **0.12** | **0.002** |
| | | June & with N | | -0.01 | 0.12 | 0.96 |
| | | August & without N | | 0.15 | 0.09 | 0.11 |
| | | August & with N | | -0.17 | 0.11 | 0.12 |
| | Sown $CWM_{SLA}$ | June & without N | All | **-0.17** | **0.06** | **0.009** |
| | Fungicide | June & without N | None | **-0.24** | **0.09** | **0.009** |
| | | June & with N | | 0.02 | 0.10 | 0.87 |
| | | August & without N | | 0.05 | 0.07 | 0.50 |
| | | August & with N | | **-0.34** | **0.10** | **< 0.001** |
| Ω_all | Sown richness | June & without N | All | **0.07** | **0.02** | **< 0.001** |
| | Sown MPD | June & without N | All | 0.03 | 0.02 | 0.17 |
| | Sown $CWM_{SLA}$ | June & without N | All | **-0.07** | **0.02** | **< 0.001** |
| | Fungicide | June & without N | All | -0.02 | 0.02 | 0.21 |
| Competitive ability | Sown | June & | All | -0.03 | 0.06 | 0.63 |

| | | | | | | |
|---|---|---|---|---|---|---|
| differences | richness | without N | | | | |
| | Sown MPD | June & without N | All | **0.24** | **0.08** | **0.002** |
| | Sown $CWM_{SLA}$ | With N | N | **-0.41** | **0.10** | **< 0.001** |
| | | Without N | | **0.20** | **0.10** | **0.04** |
| | Fungicide | June & without N | All | -0.002 | 0.06 | 0.98 |

**Table S5**. Species involved in the PaNDiv experiment. *Anthriscus sylvestris* (As) and *Heracleum sphondylium* (Hs) were not included in the phytometer experiment and were excluded from the calculation of biodiversity effects on productivity.

| Plant species (abbreviation) | Functional group | Growth form |
|---|---|---|
| *Dactylis glomerata* (Dg) | Fast growing | Grasses |
| *Holcus lanatus* (Hl) | | |
| *Lolium perenne* (Lp) | | |
| *Poa trivialis* (Pt) | | |
| *Anthoxanthum odoratum* (Ao) | Slow growing | |
| *Bromus erectus* (Be) | | |
| *Festuca rubra* (Fr) | | |
| *Helictotrichon pubescens* (Hp) | | |
| *Anthriscus sylvestris* (As) | Fast growing | Herbs |
| *Crepis biennis* (Cb) | | |
| *Galium album* (Ga) | | |
| *Heracleum sphondylium* (Hs) | | |
| *Rumex acetosa* (Ra) | | |
| *Taraxacum officinale* (To) | | |
| *Achillea millefolium* (Am) | Slow growing | |
| *Centaurea jacea* (Cj) | | |
| *Daucus carota* (Dc) | | |
| *Plantago media* (Pm) | | |
| *Prunella grandiflora* (Pg) | | |
| *Salvia pratensis* (Sp) | | |
